# NRGRank: Coarse-grained structurally-informed ultra-massive virtual screening

**DOI:** 10.1101/2025.02.17.638675

**Authors:** Thomas DesCôteaux, Olivier Mailhot, Rafael Najmanovich

**Affiliations:** Department of Pharmacology and Physiology, Université de Montréal, Montréal, QC H3C 3J7, Canada

**Keywords:** NRGRank, virtual screening, docking methods, enrichment factor (EF1), pseudo-energy interaction, molecular flexibility

## Abstract

NRGRank is a coarse-grained structurally-informed virtual screening Python package with accuracy comparable to docking-based methodologies but up to 100-fold speed increase. NRGRank is based on a coarse-grained evaluation of pairwise atom-type pseudo-energy interactions that implicitly accounts for compound and side-chain flexibility as well as limited backbone movements. We compare NRGRank to docking-based virtual screening software Glide, Autodock Vina and DOCK 3.7 on the DUD-E virtual screening benchmark using enrichment factors at 1% (EF1). We observe broad variations of EF1 values across targets, structural models and methods. For apo form or AlphaFold2 models, out of a subset of 37 targets from DUD-E, NRGRank has better EF1 values than Glide for 12 and 13 targets respectively. Even in holo form, where the accuracy of classical docking software increases, NRGRank has better EF1 values than 13, 10 and 5 targets out of 37 compared to AutoDock Vina, DOCK 3.7 and Glide respectively. Comparing the rank of true binders in Glide and NRGRank shows that true binders ranked in the top 1% are complementary between methods irrespective of the target form (AlphaFold, apo or holo). That is, utilizing NRGRank detects binders that are missed by Glide (and presumably other methods), whereas those found by Glide are missed by NRGRank. Furthermore, we observe that most hits found by NRGRank within the top 50 predictions (4.38 ± 5.49 hits on average for AF2 targets) remain once the top 1% of predictions are re-scored with Glide, but the hit rate within the top 50 predictions increases. NRGRank can evaluate one molecule in 0.3 s on average, enabling a modern laptop with 8 cores to screen 1,000,000 molecules in 24 hours – up to two orders of magnitude faster than the reported speed of DOCK 3.7, AutoDock Vina running on GPUs and Glide. NRGRank occupies a unique niche among tools for virtual screening being insensitive to structural inaccuracies but with comparable accuracy as state-of-the-art docking methods and fast as AI-based methods but without the dangers of overfitting as it is based on 780 pseudo-energy parameters. Combined with the fact that NRGRank does not require extensive or expensive computational resources or expert pre-processing of targets, it is unique in making high-performance ultra-massive virtual screening accessible to all.

## 1 Introduction

Among the different approaches in drug design, the ability to sift through large and diverse compound sets to identify potential hits offers the possibility of exploring novel areas of chemical space. Although cutting-edge robotic screening systems can evaluate an impressive 200,000 compounds per day ^1^, their high costs make them largely inaccessible and impractical for exploring the several billion compounds available today through synthesis-on-demand services. The landscape of make-on-demand chemical libraries is continuously expanding: even the restricted lead-like subset of compounds (MW *≤* 460, *−*4 *≤* SlogP *≤* 4.2, HBA *≤* 9, HBD *≤* 5, Rings *≤* 4, RotBonds *≤* 10) from the Enamine REAL library (under 10% of the entire library) boasts a staggering 10.1 billion compounds (September 2025), approximately 80% of which have a high likelihood of successful synthesis ^2^. Conducting high throughput experimental screening with a small fraction of these molecules presents significant challenges in terms of time and expense, not to mention the considerable resources required for synthesis and transportation. Virtual high-throughput screening (vHTS) offers a promising alternative, particularly for exploring novel libraries in the early stages of drug design. Such methodologies may take many forms. One common strategy involves ranking the molecules in a library through a calculated score and prioritizing those with the most favorable scores for further investigation. Studies show that enlarging the library size utilized in virtual screens can increase hit rate ^3, 4, 5^. As libraries grow, algorithms become more robust and computational power more widespread, ultra-massive vHTS (uHTS) is beginning to impact drug development, often identifying weak binding compounds ^6^ and sometimes strong binders ^7^.

In 2023, the Target Central Resource Database (TCRD) contained approximately 17,500 proteins in the human proteome for which there are no known binders and limited knowledge of protein function ^8^. Recent advances in protein structure prediction brought about by AlphaFold2 ^9^ have produced reliable structural models of all these proteins, enabling the potential use of vHTS for identifying modulators of protein function to significantly accelerate research. A recent study ^10^ compared the effect of using different types of target structures on enrichment factors at 1% (EF1) for the Glide docking method ^11^ on 37 targets from the Directory of Useful Decoys – Enhanced (DUD-E) benchmark dataset ^12^. The authors show that using an AlphaFold model or apo (unbound) structure halved the average EF1 compared to using a holo protein structure. While extra refinement steps may be taken to enhance the accuracy of virtual screening on apo or model structures, such refinement requires at the very least the knowledge of a validated binder. This information is obviously unavailable in the case of targets with no known binders. This degradation of Glide performance when moving from holo to apo/AlphaFold structure highlights its sensitivity to the quality of the target structure. Considered by many the industry standard for docking simulations, Glide outperforms all other methods using holo proteins but its performance is on par with other methods, including NRGRank as shown below, for apo proteins and AlphaFold models.

The accessibility of docking methods is often affected by the requirement of software licenses. A recent literature review ^13^ reveals that out of the 12 most widely used virtual screening software packages, only 4 are accessible for academic use free of charge: AutoDock Vina ^14, 15^, DOCK 3.7 ^16^, FRED ^17^ and LibDock ^18^. Some open-source docking methods require closed-source third-party applications, for which academic licenses are mandatory during target or ligand preparation. For instance, DOCK 3.7 relies on several third-party software components for optimal performance, such as OpenEye Omega, OEChem, JChem-base and Corina, all of which entail academic or paid licenses. These constraints heighten the barrier of entry for academic researchers to use virtual screening.

Recently, virtual screening methodologies leveraging artificial intelligence have emerged ^19^. However, these approaches come with notable drawbacks. Primarily, they tend to struggle in scenarios outside their training data. Despite exhibiting promising performance, particularly when applied to data similar to their training set^20^, they often perform poorly when confronted with unfamiliar situations ^21^. It is still unclear how well such methodologies perform, especially given the various types of bias that may be present. One example is not having a proper dataset split that includes a separate evaluation set. This can lead to optimizing parameters specifically to improve test-set performance, an instance of Goodhart’s law. Another trivial bias is analog bias. This occurs when different targets in the dataset are evolutionarily related, or when binders of the same target share common molecular scaffolds. Additionally, the Directory of Useful Decoys-Extended (DUD-E) ^12^ is widely used as a benchmark for virtual screening with physics-based docking methods. However, the DUD-E dataset embodies decoy bias due to property matching ^22^, making its use specifically for artificial intelligence methods problematic. By selecting decoys that have matched properties but are topologically different, a classifier that learns a large set of possible topological variations maintaining physical properties may effectively learn the complement of that set (true binders), i.e., the possible topologies that maintain the physical properties but are not decoys. By creating alternative decoy sets, Chen et al. demonstrated that CNN-based models achieve high accuracy due to analog or decoy biases rather than learning underlying rules that govern ligand-protein interactions ^22^. In this work, we compare NRGRank, a physics-based method, with other physics-based methods on the DUD-E benchmark dataset. Given the current absence of a widely used benchmark dataset devoid of decoy bias, we do not compare NRGRank to AI-based methods.

The surge in popularity of GPU (Graphics Processing Unit)-based software, exemplified by tools like Uni-Dock ^23^ and Vina-GPU 2.0 ^24^, is mainly driven by their remarkable speed enhancements compared to traditional CPU-based applications. For example, Uni-Dock can dock 38.2 million molecules in 12 hours, leveraging the power of 100 NVIDIA V100 GPUs. However, while the speed gains are impressive, it is essential to acknowledge the limitations of GPU-based docking software. One drawback is their reliance on technologies like CUDA, as seen in the case of the most recent version of Uni-Dock (v1.1). CUDA is a programming model exclusive to NVIDIA GPUs, which means that the software cannot be used by users lacking compatible graphics cards. This accessibility hurdle is compounded by the recent “AI boom”, which has spurred an unprecedented demand for GPUs, consequently driving up their prices significantly and creating an economic barrier to their utilization in vHTS.

Here we present NRGRank, a coarse-grained structure-informed Python package capable of screening a million compounds per day on a typical laptop CPU, requiring no manipulation of the target structure. NRGRank has complementary results and comparable accuracy when compared to existing docking methods, particularly for apo protein structures and AlphaFold models, removing barriers for ultra-massive virtual screening and opening the doors to exploring both chemical and protein space broadly and unbiasedly.

## 2 Results and Discussion

### 2.1 Performance on Apo Protein Structures and AlphaFold2 Models

Virtual screening software generally overlook the dynamic nature of side chains in favor of speed. However, changes in side-chain conformations can be crucial when working on targets with apo protein structures and/or AlphaFold2 (AF2) models. Apo proteins (without a bound ligand) can be structurally distant from the holo (ligand-bound) form: the binding pocket may not be well-defined or may undergo conformational changes upon ligand binding. In 90% of binding-sites, at least one side-chain undergoes a rotamer change upon ligand binding and in approximately a third of cases the rotamer change is critical for ligand binding ^25^. AF2 provides accurate predicted protein structures, but these predictions cannot account for the full range of native protein conformational flexibility. Predicted models may have inaccurate positions for some side-chains, or may reflect a local minima incompatible with ligand binding. In virtual screening scenarios, traditional rigid-protein docking methods do not handle these cases well. To address this limitation, NRGRank adopts a coarse-grained methodology that implicitly accounts for side chain and ligand flexibility. NRGRank calculates binding energy on a per index-cube basis rather than a fixed distance. That is to say, the contribution to the overall interaction energy of a ligand atom within an index-cube interacting with a protein atom in an adjacent index-cube is the same irrespective of the specific locations of each atom within their respective index cubes. We experimented with a wide range of index-cube length values (Supplementary Figure S1), as well as varying the number of conformers per ligand (Supplementary Figure S2) and the number of rotations per axis (Supplementary Figure S3) on the DUD-E dataset. Despite these variations, EF1 results were largely insensitive to changes in these parameters when they remained near the default values: a single conformer per ligand, 9 rotations per axis (729 in total), and an index-cube length of 6.56 Å (see Methods). This suggests that most ligand conformational changes, side-chain rotamer changes, or minor backbone movements likely keep the relevant atoms within the same index-cube. As a result, the method is less sensitive to the effects of dynamics in ligand binding, while also simplifying and accelerating the algorithm.

We use the DUD-E dataset to compare the performance of NRGRank against that of Glide for apo protein structures and AF2 models. The comparison is restricted to Glide as we are not aware of detailed (per ligand/target pair) published data for apo targets or AF2 models for DOCK 3.7 or AutoDock Vina. The 37 targets selected by Zhang *et al.* to evaluate Glide ^10^ are cases for which apo and holo structures are provided in DUD-E. The results show considerable variability in EF1 for both Glide and NRGRank when analyzing apo and AF2 structures for all targets. Specifically, the average EF1 value for NRGRank with apo structures is 5.4 ± 5.7, while for Glide with apo structures the EF1 average is 10.4 ± 8.1. For AF2 structures, NRGRank yields an average EF1 value of 5.4 ± 6.0, while Glide has a value of 11.0 ± 10.9. A paired Student’s t-test between NRGRank and Glide yields p-values of 0.003 for apo structures and 0.004 for AF2 structures. The EF1 values for individual apo and AF2 targets are shown in Figure 1. Despite the average EF1 differences, NRGRank has higher individual EF1 values than Glide for 14 targets (representing just over a third of this DUD-E subset) for apo form and AF2 models (Figures 1B and 1C). The individual values of EF1 for each target in the different forms for each software can be found in Supplementary Table S1 (available in the Zenodo data depository).

**Figure 1:**
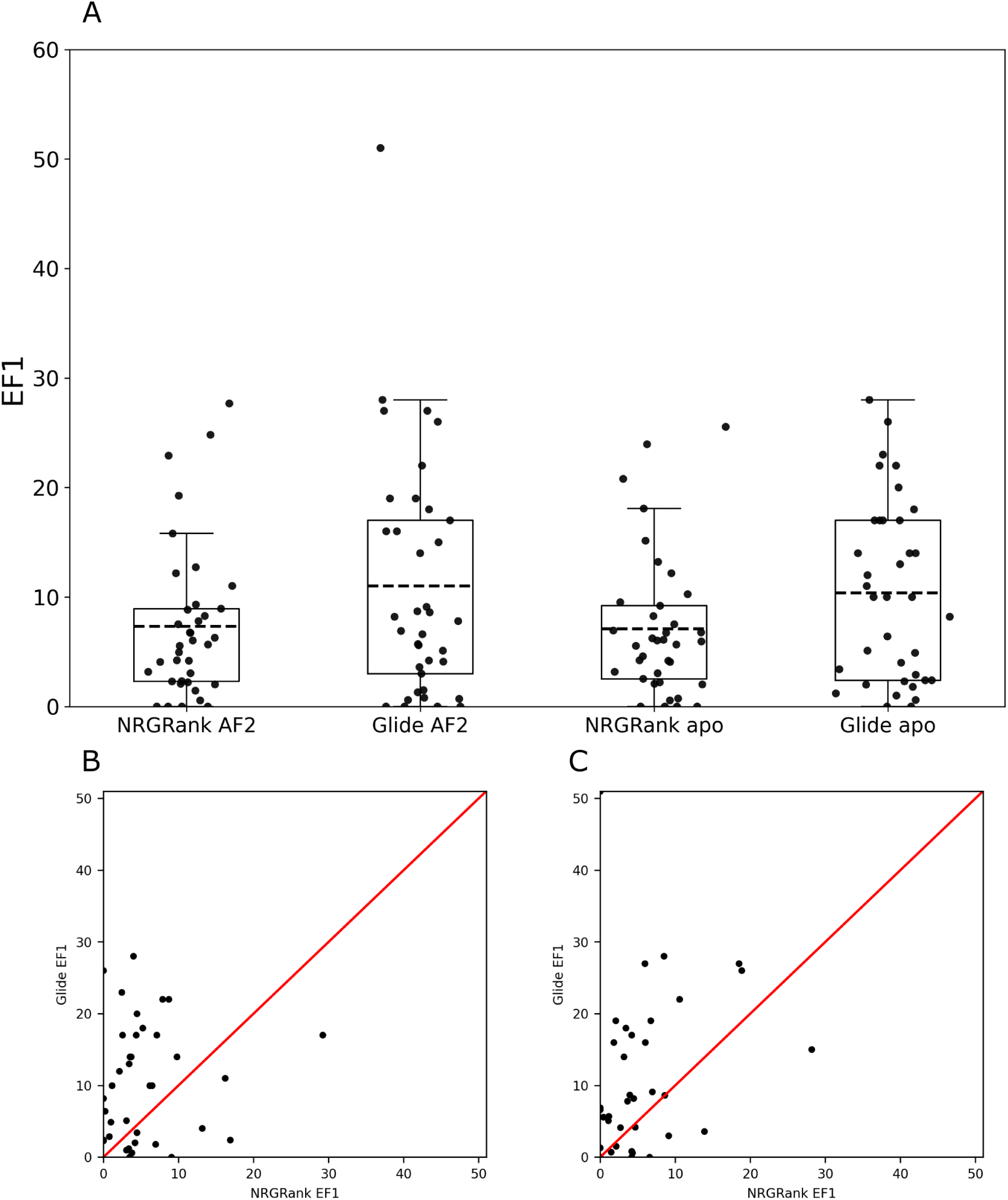
Enrichment factors for non-holo targets. EF1 for the 37 DUD-E targets with published data ^10^ for AlphaFold2 models (AF2) and apo form targets for Glide and NRGRank (A). Comparison of EF1 for individual targets shows that NRGRank has better EF1 values for 12 targets in apo form (B) and 13 targets for AF2 models (C) out of 37. Across the full set of targets, paired *t*-tests indicate that NRGRank and Glide EF1 values are significantly different for both apo (*p* = 0.0035) and AF2 (*p* = 0.0040) cases.

**Table 1:** Average EF1 values for different software on the subset of 37 DUD-E targets by target structure.

|  | Target type<sup>a</sup> |  |  |
| --- | --- | --- | --- |
|  | AF2 | Apo | Holo |
| **NRGRank** | 5.4 ± 6.0 | 5.4 ± 5.7 | 6.9 ± 8.0 |
| **AutoDock Vina** | NA | NA | 10.0 ± 8.6 |
| **DOCK 3.7** | NA | NA | 14.2 ± 10.8 |
| **Glide** | 11.0 ± 10.9 | 10.4 ± 8.1 | 22.4 ± 13.2 |

Despite the differences in performance between NRGRank and Glide for apo or AF2 models, each software identifies distinct subsets of true binders. Figure 2 shows the rank of individual true binders across all 37 DUD-E targets for which we can compare our results with those of Glide for AF2, apo and holo targets. There is a high number of ligands with rank above 1% for both NRGRank and Glide (quadrant 1 in all panels) representing true positive binders missed by both software. However, there are large numbers of molecules that have a rank below 1% for one software but not for the other (quadrants 2 and 4 in all panels). Only a very small fraction of true positives are detected by both software within the top 1% (quadrant 3 in all panels). While intuitively users of virtual screening docking software would be tempted to use multiple software and focus on the intersection of top results as independent validation (quadrant 3), our results indicate that with this strategy most true binders for any target would be discarded. Therefore, despite differences in EF1 values, NRGRank detects true binders that are missed by other software.

**Figure 2:**
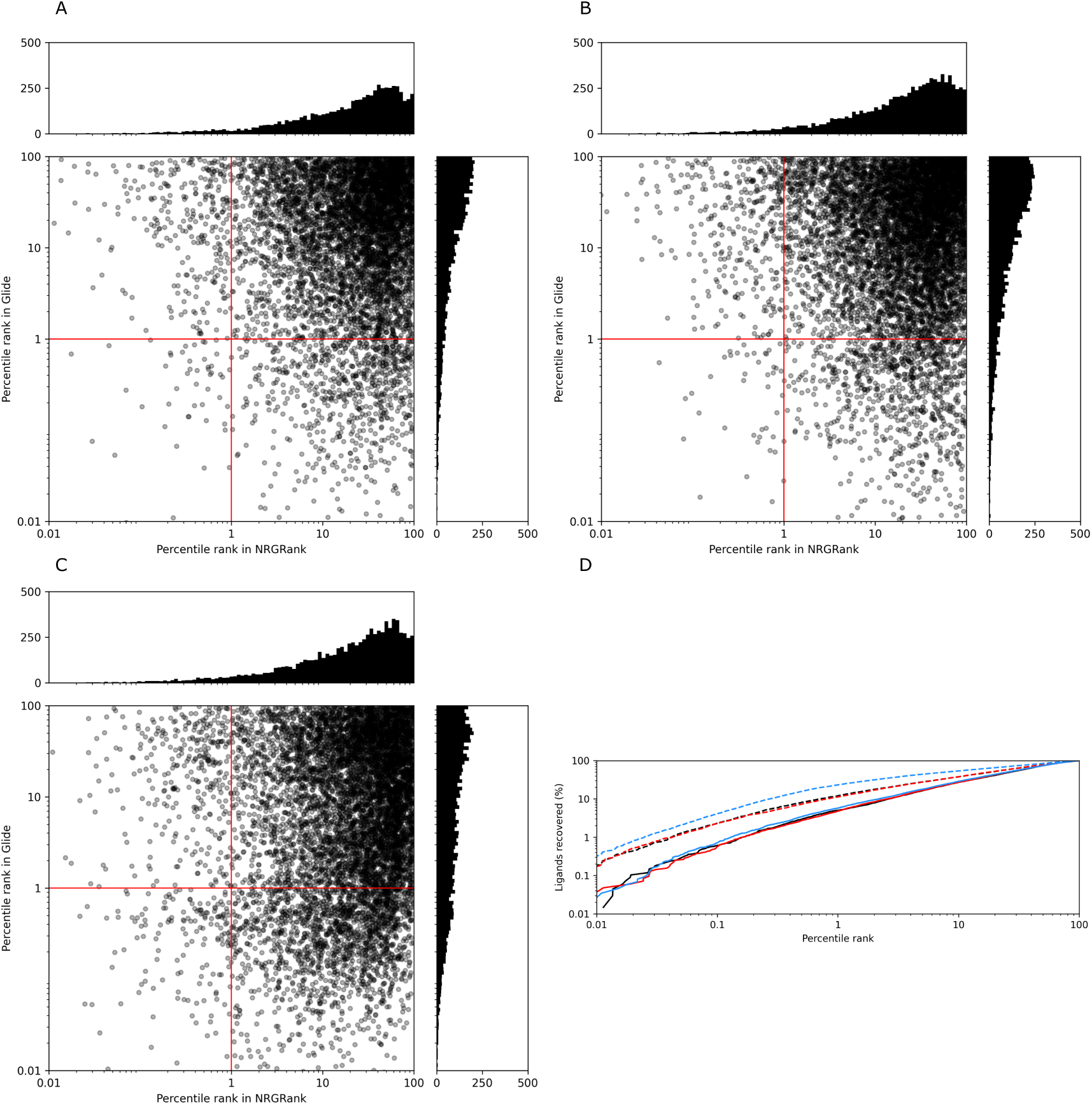
Comparison of true binder ranking in NRGRank and Glide across 37 AF2 (A), apo (B), and holo (C) targets. The number of true binders missed by both methods (quadrant 1) was 5,475, 6,856, and 6,164 for AF2, apo, and holo targets, respectively. Binders identified by NRGRank but missed by Glide (quadrant 2) numbered 289, 352, and 337 for AF2, apo, and holo targets, respectively. Conversely, binders missed by NRGRank but identified by Glide (quadrant 4) totaled 723, 886, and 1,087 for AF2, apo, and holo targets, respectively. Binders identified by both methods (quadrant 3) were 50, 42, and 161 for AF2, apo, and holo targets, respectively. (D) Cumulative distributions of true binder ranks for NRGRank (solid lines) and Glide (dashed lines) across AF2 (black), apo (red), and holo (blue) targets. Curves that rise more steeply indicate better performance, as a greater fraction of true binders are recovered at higher ranks.

### 2.2 PERFORMANCE ON HOLO PROTEIN STRUCTURES

When the use case scenario is the discovery of novel chemical matter for undrugged targets or cryptic binding pockets, apo forms and modeled structures serve as the most faithful benchmark for virtual screening performance, hence the previous section. However, docking software is often evaluated using holo protein structures ^26^, in part because there are several legitimate applications of molecular docking where a holo structure is available as a starting point, but also because side-chain flexibility is rarely modelled, hindering the performance on apo structures. Nevertheless, we use holo form structures provided with the DUD-E dataset to compare NRGRank to widely used docking software DOCK 3.7, AutoDock Vina and Glide. As data for Glide was only available for the subset of 37 targets selected by the authors who analyzed Glide ^10^, we restricted our analysis to those 37 targets. NRGRank achieves an average EF1 of 6.9 ± 8.0 for holo targets while Glide obtains an average EF1 of 22.4 ± 13.2. The same situation observed for Apo and AF2 targets is true here again, namely, when comparing the ranks of individual binders, true binders detected by NRGRank within the top 1% are largely missed by Glide and those found by Glide among the top 1% predictions are missed by NRGRank. Therefore, despite a lower EF1 value for holo protein structures (which was expected since holo structures are an ideal use case of Glide), it is still true that NRGRank is complementary to Glide (Figure 2C). On this same subset of DUD-E, the average EF1 values for AutoDock Vina and DOCK 3.7 are respectively 10.0 ± 8.6 and 14.2 ± 10.8. Despite mean EF1 values for holo structures being smaller for NRGRank, the variance for all software is high (Figure 3A). Comparing the EF1 of individual targets between NRGRank and other software for holo target structures (Figure 3B and 3C) shows that NRGRank has higher or equal EF1 values for 13 targets relative to AutoDock Vina and 10 relative to DOCK 3.7. Despite the 3-fold difference in average EF1 values with respect to Glide, for 5 targets, NRGRank still has better EF1 values than Glide (Figure 3D).

**Figure 3:**
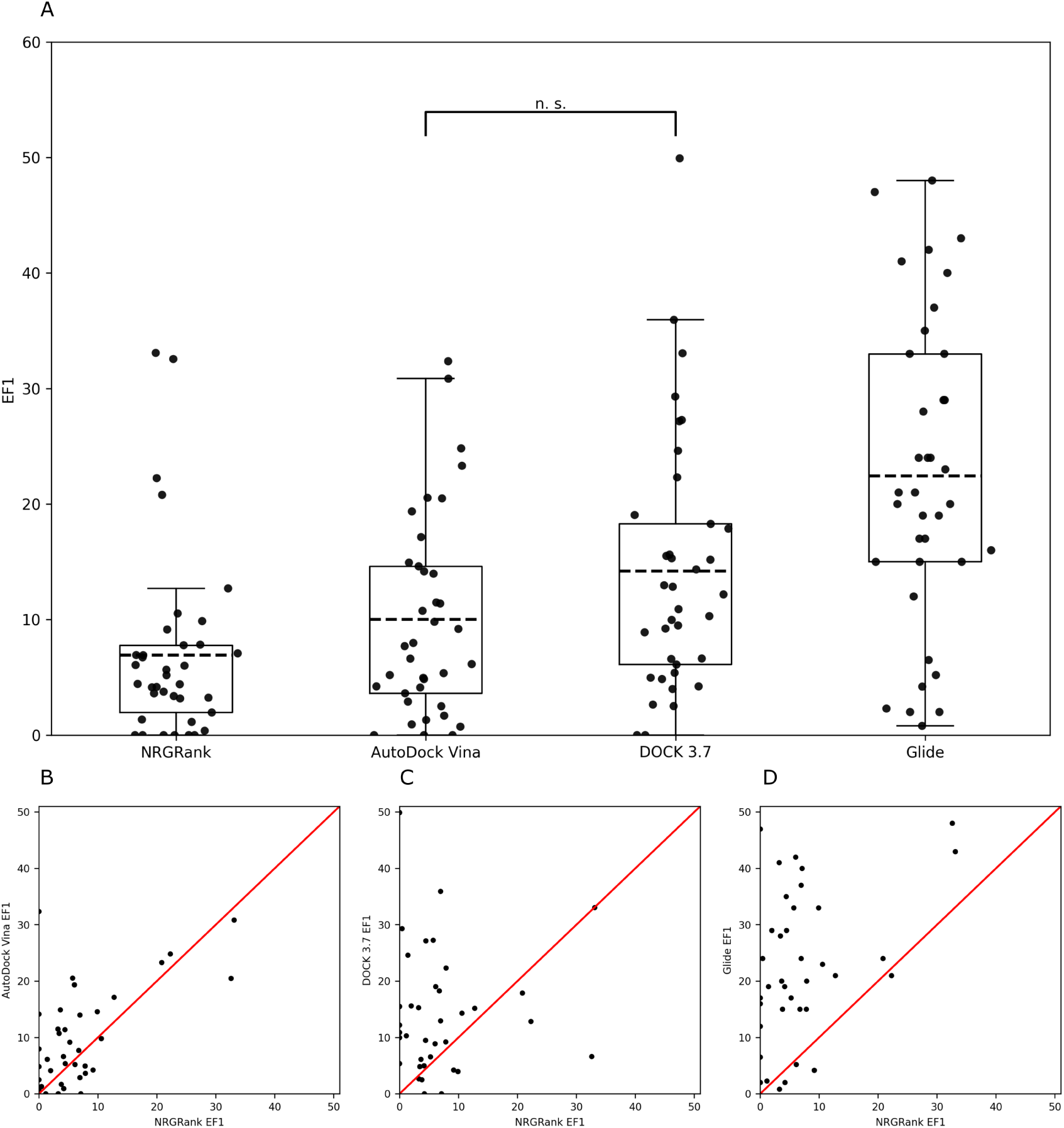
Comparison of enrichment factors (EF1) for NRGRank and other docking methods across 37 DUD-E targets in holo form. (A) Boxplots showing EF1 distributions for NRGRank, AutoDock Vina, DOCK 3.7, and Glide. Each point represents one target. Pairwise Student’s t-tests showed significant differences for NRGRank vs. DOCK (p = 0.0024), NRGRank vs. Glide (p < 0.0001), and Glide vs. DOCK (p = 0.0051). The EF1 differences between Vina and DOCK (p = 0.0745) was not significant. (B–D) Scatter plots comparing EF1 values of NRGRank with AutoDock Vina (B), DOCK 3.7 (C), and Glide (D). The red diagonal line indicates equal performance. Overall, out of the 37 targets, NRGRank outperforms AutoDock Vina, DOCK 3.7, and Glide for 13, 10, and 5 targets, respectively.

### 2.3 RE-SCORING NRGRANK RESULTS WITH GLIDE

While NRGRank is a method for ranking small-molecules in virtual screening, unlike docking methods, it is not designed to predict the correct ligand pose and generate a ligand-protein complex structure. In situations where a docked pose in virtual screening is required, we envision the use of a docking software (such as AutoDock Vina, DOCK 3.7, Glide or FlexAID among others) to be used to re-score the top fraction of the NRGRank output. In order to simulate such a situation with Glide, we re-scored the top 1% predictions of DUD-E compounds (including both true binders and decoys) for a given target in the rank order defined by their Glide scores. The relative position in the ranked list for each target of the true binders represents the effect of docking such molecules with Glide. In practical terms we decided to restrict the analysis to targets with at least 50 true binders and calculate how many true binders are among the top 50 predicted molecules before and after re-scoring with Glide. This experiment reflects a real scenario in which one would use NRGRank to rank a large dataset, re-score a small fraction of top ranked molecules and have the top 50 compounds synthesized for experimental validation. The value 50 for the number of molecules to be tested experimentally was chosen here to balance having a number that would be reasonable to test experimentally and the number of targets in the subset of DUD-E for which Glide scores are available. This experiment was performed using AF2 models, as well as apo and holo protein forms for 34 targets (marked in Supplementary Table S1). For AF2 models, a total of 329 binders were among the 1% NRGRank predictions with 117 among the top 50 predictions for all 34 targets (quadrants 2 and 3 in Figure 4A). Of these 117, 111 remain among the top 50 predictions even after re-scoring (quadrant 3 in Figure 4A), the remaining 6 are lost with re-scoring (quadrant 2 in Figure 4A) and an additional 69 binders that had ranks above 50 with NRGRank are added to the top 50 list after re-soring (quadrant 4 in Figure 4A). Re-scoring increases the fraction of binders in the top 50 from 45% to 64% (Figure 4B shows the hit rate distribution). For apo targets (Figures 4C and 4D), we have 145 in quadrant 1 (missed with or without re-scoring), 147 in quadrant 3 (in the top 50 before and after re-scoring), 11 in quadrant 2 (lost upon re-scoring) and 84 in quadrant 4 (added to the top 50 upon re-scoring). The fraction of binders in the top 50 increases from 40% to 59%. For holo targets (Figures 4E and 4F), we observe 135, 11, 217 and 122 in quadrants 1, 2, 3 and 4 respectively with a fraction of binders in the top 50 increasing from 47% to 90% upon re-scoring. While re-scoring is beneficial, it is by no means essential in order to find binders among the top 50 predictions. In 28 out of the 34 targets, we find at least one binder among the top 50 predictions. The hit rate distributions show that for all three types of structures (AF2, apo, holo) the re-scoring shifts the distributions right, increasing the hit rate for the 34 targets (Figure 4B, D & F).

**Figure 4:**
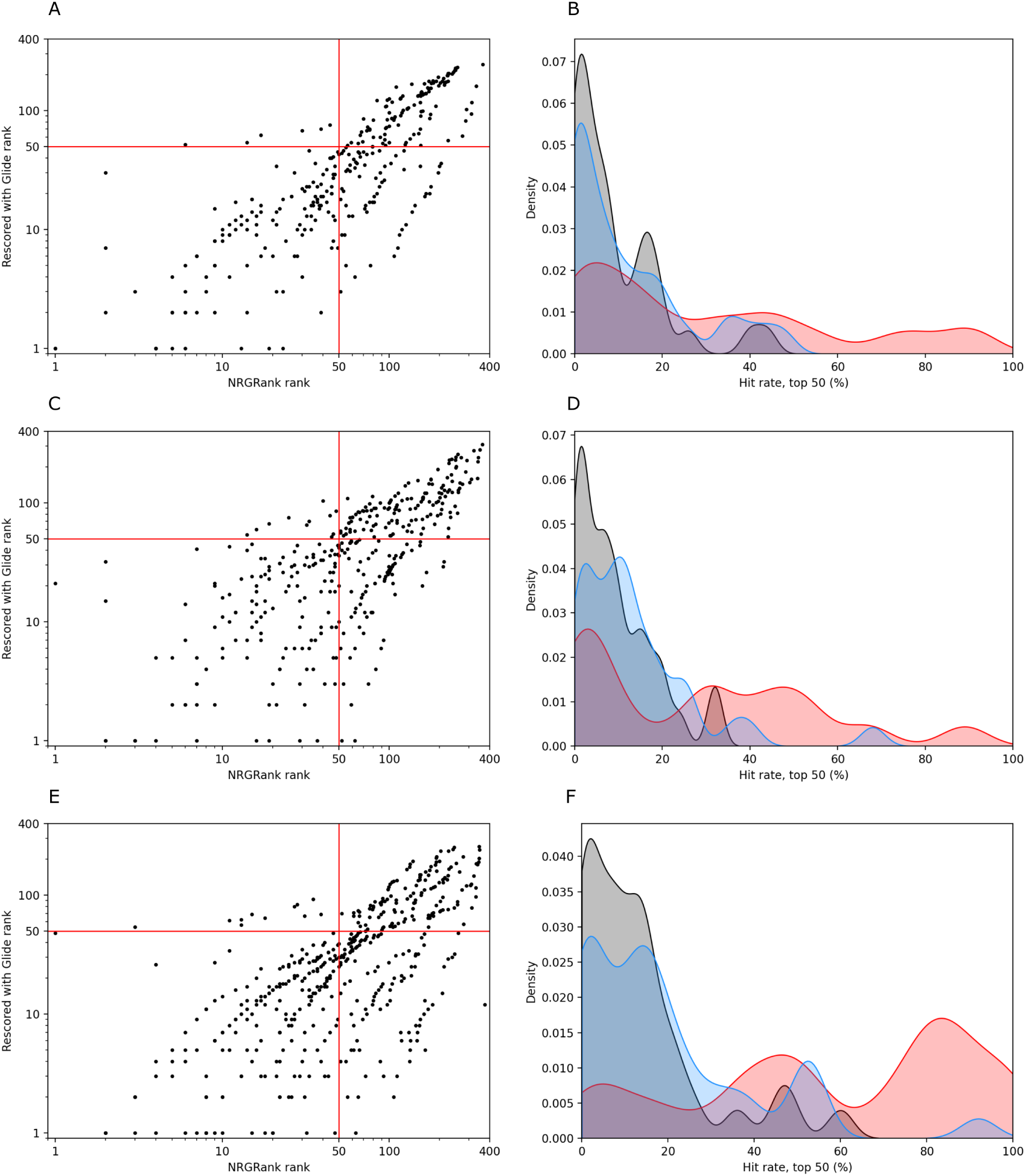
Effect of re-scoring NRGRank hits with Glide. For each of 32 targets in DUD-E with at least 50 true binders, we re-scored the top 1% ranked molecules (both binders and decoys) by re-ordering the molecules in the relative order they appear in the published Glide results and determined how many molecules remain, are added or removed from the top 50. Quadrant 1 in panels A (AF2), C (apo)& E (holo) represent binders with rank above 50 before or after re-scoring. Quadrant 3 represent molecules that were in the top 50 predictions and remain after re-scoring. Quadrant 2 represents binders removed from the top 50 upon re-scoring. Lastly, in quadrant 4, are binders that were added to the top 50 upon re-scoring. Panels B (AF2), D (apo) & F (holo) show the hit rate distributions with black, red and blue corresponding to NRGRank, Glide and NRGRank rescored with Glide respectively.

### 2.4 EVALUATION OF NON-PROPERTY MATCHED DECOYS

By virtue of the high level of similarity between known binders and decoys (assumed to be non-binders), the DUD-E dataset defines a lower bound on the accuracy of any biophysics-based vHTS method (not so for artificial intelligence systems as mentioned in the introduction). To simulate a more common scenario, we assessed NRGRank’s ability to distinguish true binders from non-property-matched molecules. The Enamine REAL diversity subset (ERDS) compound library, representing 1% of the entire REAL dataset, was used for this experiment. At the time of download the library contained 67.5 million compounds (available for download from our Zenodo repository for reproducibility purposes), while the equivalent dataset available today from Enamine contains around 95.6 million compounds. The true binders for each target in DUD-E were selected from ChEMBL as molecules matching any of the following criteria: IC50, EC50, K*_i_* or K*_d_* values *≤* 1 *µ*M. This is the same criteria used to select binders by the original DUD-E authors. We plot in Figure 5A for each of the 37 targets selected by Zhang *et al.* the mean z-score and standard deviation of known binders calculated from the mean and standard deviation of ERDS docking scores. The mean z-score for known binders is negative for all 37 targets and well below zero for most targets, with an average mean z-score of -1.87. When expanding the analysis to all 102 DUD-E targets, all targets except for 4 have a negative mean z-score (Supplementary Figure S4A) and the average mean z-score is -1.66. This result shows that NRGRank separates binders from random molecules. Considering the overlap between the distributions (Figure 5B and Supplementary Figure S4B), for all targets there are large numbers of ERDS molecules with ranks better than the rank of known binders. For 67 targets, some ERDS molecules have better scores than known binders. For example, 23 ERDS molecules rank better than the best binder for the *β*1-adrenergic G-protein coupled receptor (DUD-E ID adrb1). This behavior is expected at the scale of screening performed here: the ERDS library has 4 to 6 orders of magnitudes more compounds than the known binders for the targets.

**Figure 5:**
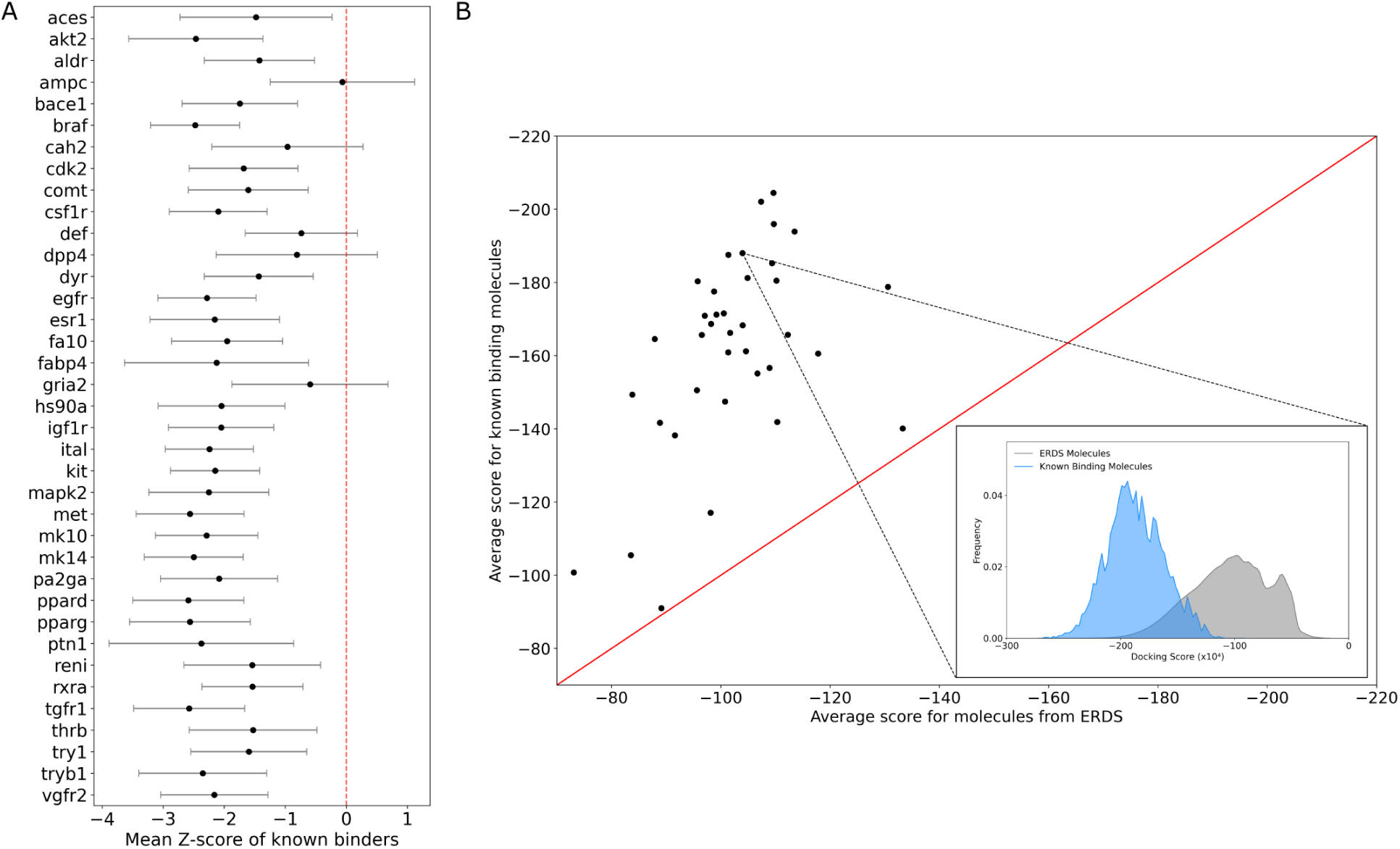
Screening ERDS (Enamine REAL Diversity subset) on 37 targets of the DUD-E dataset. (A) Mean Z-score ± standard deviation for known binders of each target, calculated from the mean and standard deviation of ERDS molecule docking scores. The dashed red line at zero indicates the expected average for random compounds. (B) Comparison of average ERDS scores for known binders (y-axis) versus ERDS compounds (x-axis) across targets. Points above the diagonal line correspond to targets where known binders score better than ERDS molecules. The inset shows the distribution of docking scores for the BRAF protein kinase, with 4,048 known binders in DUD-E (blue) contrasted against ERDS molecules (gray).The list of ERDS and known binder distribution averages as well as number of known binders can be found in Supplementary Table 2.

### 2.5 Finding unique molecules for each target

While NRGRank achieves promising retrospective performance across DUD-E, we wondered whether its coarse-grained scoring function could be subject to artifacts when used for prospective ultra-large library screening. Indeed, it could be that some molecules score invariably well across all targets, acting as vPAINS (virtual PAINS ^27^ or **<u>P</u>**an-**<u>A</u>**ssay **<u>IN</u>**terference compound**<u>S</u>**). To our knowledge, this is the first time vPAINS are discussed in virtual screening. However, it stands to reason that scoring functions have biases that may be reflected in top scoring compounds. It is our hope that the assessment of vPAINS compounds becomes current for virtual screening methods. To investigate this question in the context of NRGRank, we computed the pProp metric for every molecule from the ERDS set, against each of the 102 targets in the full DUD-E set. The pProp value is defined as the negative logarithm of the fractional rank of a molecule ^5^. For instance, a pProp of 2 corresponds to top 1%, a pProp of 3 to top 0.1%, and so on. Here, we also define ΔpProp: the difference between the pProp of a molecule for a given target minus its median pProp across all 102 DUD-E targets. A molecule is considered an interesting candidate for a target if it has a pProp *≥* 3 (corresponding to the top 0.1% of ranked compounds) and a ΔpProp > 2, indicating strong enrichment for a specific target relative to the background. An average of 811 molecules per target fit these criteria and all targets have at least 10 interesting candidates (Figure 9), with adenosylhomocysteinase (refered as sahh in DUD-E) standing out with 47,778 candidate molecules. Even under a more stringent ΔpProp threshold of 3 and pProp threshold of 4, 89/102 targets still retained at least one interesting candidate and an average of 13 molecules per target fit these criteria. These results suggest that while vPAINS compounds are present in NRGRank (molecules with high pProp but low ΔpProp), as they probably are with all virtual screening software ^4^, target-specific compounds are always found in the top-scoring region of an NRGRank screen. Furthermore, the use of pProp and ΔpProp can make their detection possible. Given NRGRank’s speed, a similar approach could be used in prospective scenarios, where one could score top compounds across hundreds of cross-targets and only keep candidates with high predicted selectivity for the target under study.

### 2.6 The Good, the Bad and the Ugly: Three practical examples of top hits

#### The Good

We filtered the top-scoring compounds across all DUD-E targets by selecting molecules with pProp *≥* 4 and ΔpProp *≥* 3. This corresponds to molecules ranked within the top 0.01% of ERDS scores (rank 6750 or better) and at least three orders of magnitude better ranked for a given target compared to their median rank across all 102 targets. After filtering, the target with the largest number of retained molecules was sahh (445 compounds). In DUD-E, sahh is represented by PDB structure 1LI4, which corresponds to the human S-adenosylhomocysteine hydrolase bound to the high-affinity inhibitor Neplanocin A (PDB ligand code: NOC), with an experimentally measured *K_i_* of 3.8 nM against bovine liver sahh ^28^.

We then applied our docking software FlexAID ^29^ (2000 generations and chromosomes for a total of 4*x*10^6^ energy evaluations per molecule) to all 445 molecules and selected promising candidates based on several criteria: FlexAID score, structural dissimilarity from NOC (Tanimoto coefficient below 0.5), cLogP, and the presence of functional groups of interest. One candidate in particular stood out: s_994 (Figure 6). Many of the key interactions observed between NOC and sahh in 1LI4 ^30^ are also present in our model of S_994 bound to sahh (Figure 6). We re-scored S_994 across all 102 targets using FlexAID and observed a Pearson correlation coefficient of 0.65 between NRGRank scores and FlexAID scores (Supplementary Figure S5). Based on our experience, FlexAID scores below *−*200 AU are generally indicative of successful binding. For S_994, the best FlexAID score was *−*246.49 AU. Across all 102 DUD-E targets, the mean FlexAID score for S_994 was *−*215.75 *±* 41.61 AU.

**Figure 6:**
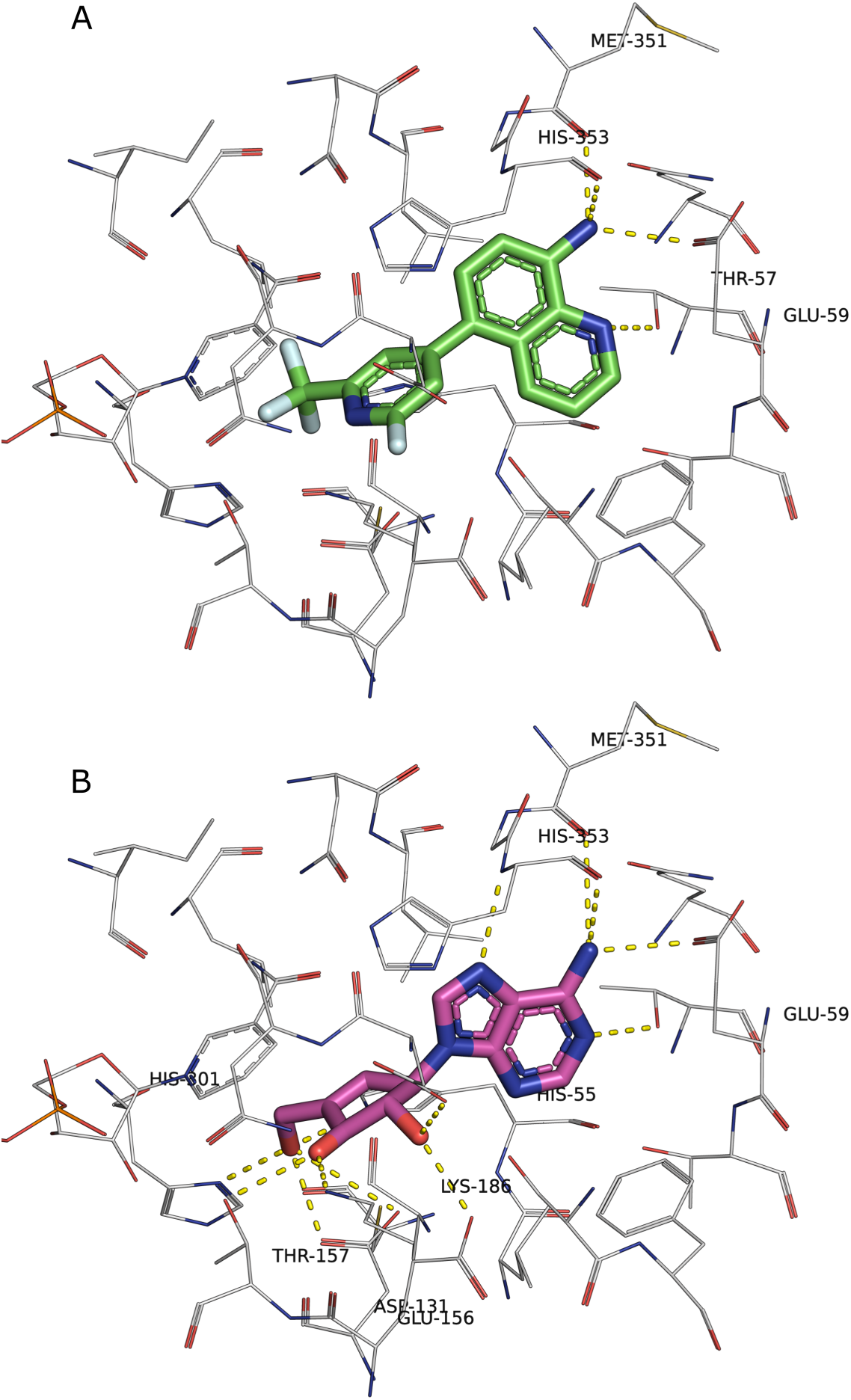
FlexAID docking pose of ERDS compound *S*_994 on sahh. The ERDS compound *s*_271570 19619370 24431994 in green (S_994), was docked with FlexAID using 2000 generations and chromosomes. One of the top scoring pose shares several moieties with the compound NOC (Neplanocin A in magenta) in the PDB structure 1LI4 used as a target. For S_994, the best FlexAID score was *−*246.49 AU, while the pose shown here scored *−*62.27 AU. We selected this pose despite its higher CF because it highlights a known H-bond interaction between GLU59 and Neplanocin A.

#### The Bad

The PDB structure assigned to adrb1 by DUD-E is 2VT4. The receptor is the turkey *β*1-adrenergic receptor (adrb1) bound to the high affinity antagonist cyanopindolol (pdb ligand code P32). The affinity of this molecule for the guinea pig adrb1 ranges from 27 to 40 pM ^31^, equivalent to *−*14.30 to *−*14.18 kcal/mol. We then proceeded to dock the top-ranking ERDS compound (Enamine ID s_22 22609224 10640652 or S_652 for short) with our software FlexAID ^29^. Many of the relevant interactions observed between P32 and adrb1 in 2VT4 ^32^ are equally present in our model for S_652 interacting with adrb1 (Supplementary Figure 6). We re-scored molecule S_652 on all 102 targets with FlexAID and find a Pearson’s correlation coefficient between the NRGRank scores and FlexAID scores of 0.62. The FlexAID score for S_652 is -258.03 AU. The median FlexAID score for S_652 on all 102 DUD-E targets is *−*87.73. At face value, this result would indicate an interesting candidate. However, S_652 is among the top 100 ranked molecules for 40 targets. This may hint at this molecule being an artifact of the method. However, it is important to note that only 12 targets have FlexAID scores below -200, including among them adrb2 (also known to tightly bind cyanopindolol). Of these, there are only 3 DUD-E targets with better FlexAID scores than adrb1 (glcm, cp2c9 and ppara with scores of *−*269.63 AU, *−*263.27 AU and *−*259.13 AU respectively). Whereas the NRGRank ranks point to S_652 as a potential artifact, the FlexAID scores do not. The fact that S_652 ranks among the top 100 molecules for 40 DUD-E targets suggests that the scoring function in NRGRank captures signals beyond what makes the binding-sites of these diverse targets different. One possibility is a signal from backbone atoms. However, a version of NRGRank only considering interactions with protein side-chain atoms has lower performance (average EF1) on the DUD-E than the version presented here (data not shown).

#### The Ugly

Certain molecules exploit biases in the scoring function and consistently obtain favorable docking scores across most targets in the DUD-E dataset. We first examined which ERDS compounds appeared most frequently among the top 1000 ranked molecules for each target. One striking example is Enamine ID s_527 9154220 21069240 (hereafter referred to as S_527). This molecule appears in the top 1000 rankings for 77 out of 102 targets, with a mean pProp of 5.52 and a median pProp of 5.60 (corresponding to a mean rank of 1643 and a median rank of 171). When rescored with FlexAID across all 102 targets (using 2000 generations and chromosomes), S_527 achieved an average CF of *−*182 AU, with CF values below *−*200 for 40 targets. Importantly, S_527 has a mean ΔpProp of only 0.01 across all targets, meaning it would be eliminated by applying a ΔpProp *>* 2 filter. Considering the discussion above, we believe it is beneficial to subject top-ranking compounds to additional scrutiny including docking against a number of unrelated targets to be able to calculate a Δ, as is the case with S_652 on adrb1 and S_527, and determine if the compound ranks highly on multiple unrelated targets.

### 2.7 ANALYSIS OF TOP SCORING COMPOUNDS ON CLOSELY RELATED TARGETS

The capability of binding a particular target without binding similar, closely-related targets is a desired objective in drug design. As NRGRank’s coarse-graining ‘bundles’ multiple atoms in index-cubes and sums over 27 index-cubes, it could render the overall contribution of small binding site differences to scoring insignificant and make it impossible to identify selective compounds for similar binding-sites. To investigate whether it is possible to find unique top scoring molecules for closely related proteins, we screened the ERDS compound library of 67.5M molecules on three closely related kinases.

The proto-oncogene serine/threonine kinase PIM1 is considered as a potential target in multiple types of cancer including prostate and triple negative breast cancer ^33, 34, 35, 36, 37, 38, 39^. Two other kinases PIM2 and PIM3 belong to the same family. There are six differences between residues lining the PIM1, PIM2 and PIM3 binding-sites (Figure 7). However, the side-chains of four among these are facing away from the cavity. Of the remaining two positions with cavity-facing side-chains among the three PIM kinases, there are two differences between PIM1 and PIM2: VAL126^PIM1^ALA122^PIM2^ and GLU124^PIM1^-LEU120^PIM2^, while between PIM1 and PIM3 only one: VAL126^PIM1^-ALA129^PIM3^, and between PIM2 and PIM3 also only one difference: LEU120^PIM2^-GLU127^PIM3^ (Figure 7).

**Figure 7:**
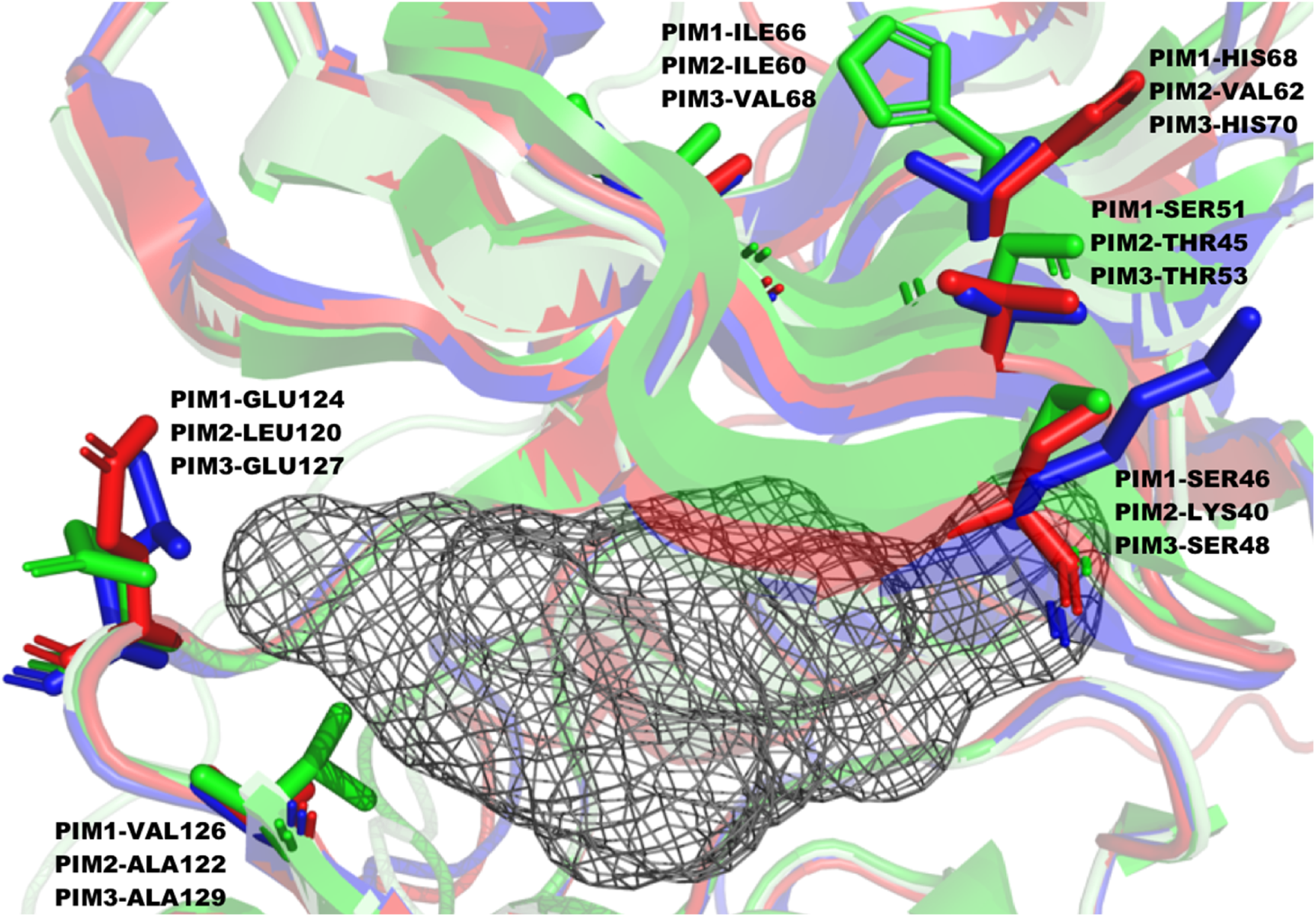
Binding-site differences between PIM1, PIM2 and PIM3. The C-terminal domain is depicted at the top. There are 6 positions with differences between PIM1 (green), PIM2 (blue) and PIM3 (red) on residues lining the ATP binding-site cavity (gray). The four mutated residues on the C-terminal domain have their side-chains facing away from the ATP binding-site. Only the two mutated positions on the hinge region between the N and C-terminal domains, have their side chains facing the cavity. Of these, only one position have a change in amino-acid type. The second position maintains a hydrophobic residue. The structures used for PIM1, 2 and 3 are 1XWS, AF-Q9P1W9-F1-model_v4 and AF-Q86V86-F1-v4 respectively.

Only four molecules within the top 100 ranked molecules for PIM1, are also among the top 100 ranked molecules for either PIM2 or PIM3 with a significant number ranked 10,000 or higher in either (Figure 8). Similarly, there are several top-ranking molecules for PIM2 (Supplementary Figure S7) and PIM3 (Supplementary Figure S8) that rank more poorly on the other members of the family. Furthermore, the top 100 ranked molecules for PIM2 have better ranks on PIM3 than for PIM1 (most molecules lie under the diagonal in Supplementary Figure S7) in agreement with the smaller number of mutations between 2 and 3, particularly considering the residues with side-chains facing the cavity. The same is true for top ranking PIM3 molecules, they rank better in PIM2 than in PIM1 (Supplementary Figure S8), but in this case, the molecules have better ranks for PIM1 (the points are shifted left) in agreement with the fact that there are less mutations between PIM3 and PIM1 versus PIM2. These results suggest that despite its coarse-grained scoring functon, NRGRank can find target-specific compounds even when presented with closely-related binding sites.

**Figure 8:**
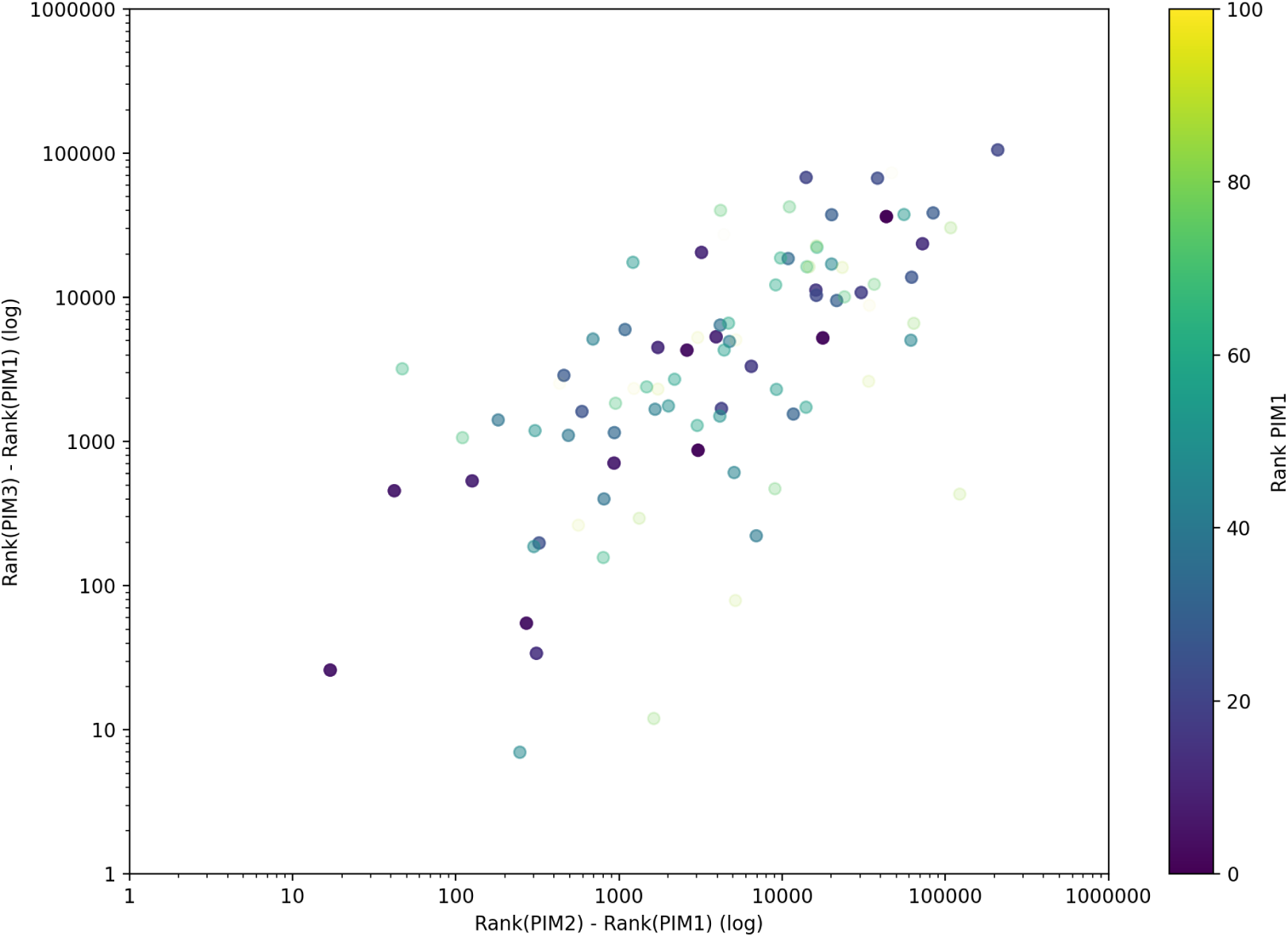
Rank comparison on PIM2 and PIM3 for the top 100 molecules ranked for PIM1. Among molecules in the top 100, a large number have poor ranks (10,000 and above) on PIM2 and PIM3.

**Figure 9:**
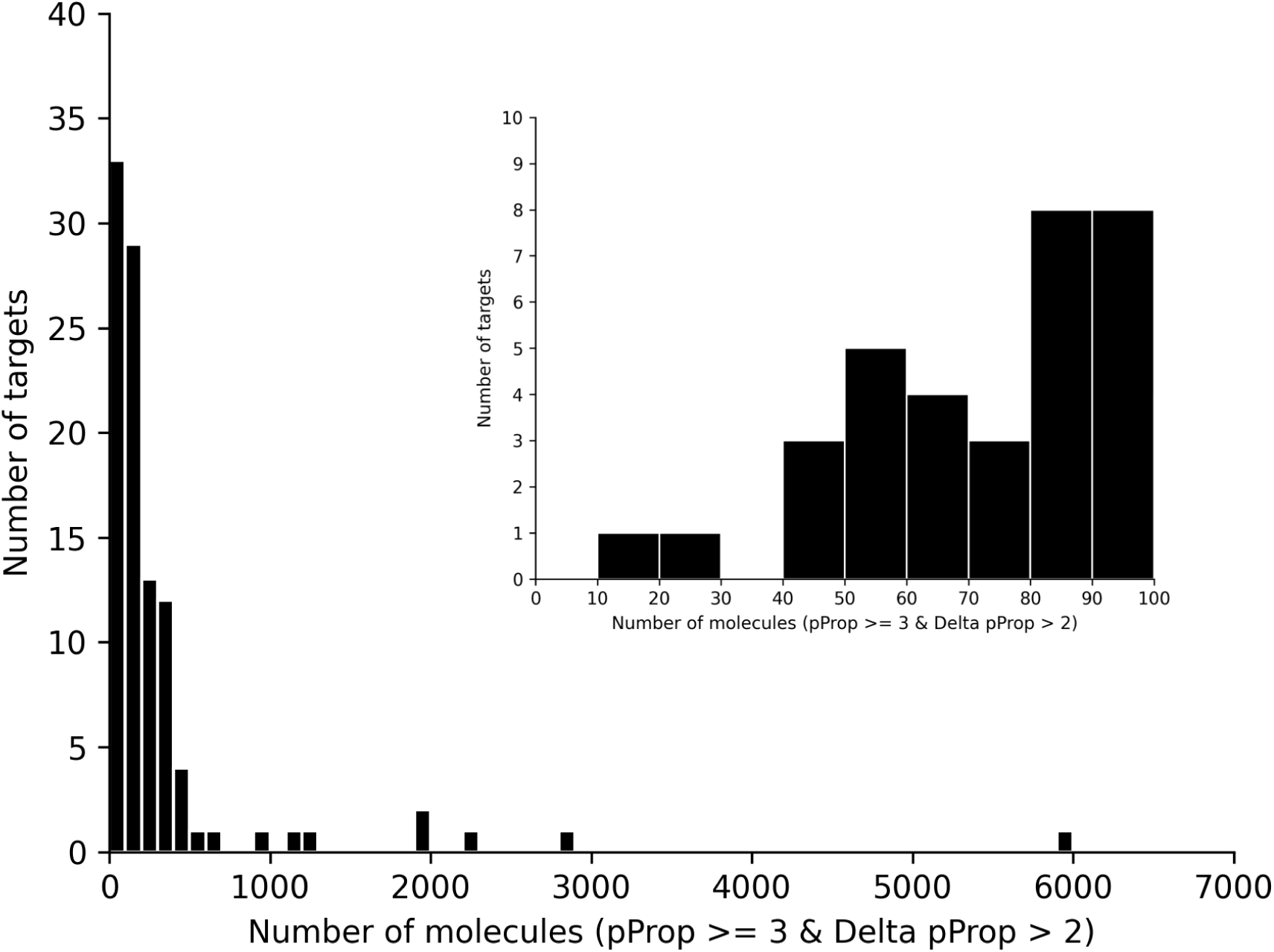
Distribution of interesting candidate molecules across DUD-E targets. Most targets have relatively few molecules meeting the pProp *≥* 3 and ΔpProp > 2 criteria, while a few contain thousands. The inset zooms in on the 0–100 range, revealing that all targets have at least 10 candidates. One extreme outlier (sahh) with 47,778 candidates is excluded for clarity. The data used is available in Supplementary Table S3

### 2.8 SPEED AND PERFORMANCE

NRGRank needs 0.36 *±* 0.23 s seconds on average to screen one molecule when testing the whole DUD-E dataset. This speed comes as a result of the pre-processing, the simplicity of the scoring function that unlike the original FlexAID scoring function ^29^ does not necessitate the calculation of surface areas in contact and lastly, due to the fact that for a given ligand atom, all interacting protein atoms within the 26 cubes of 6.56 Å plus the cube containing the ligand atom contribute equally to the docking score. This makes it possible to account implicitly for different ligand conformations.

### 2.9 DATA AND CODE AVAILABILITY

NRGRank can installed with pip and the source code is available on our GitHub repository: https://www.github.com/NRGlab/NRGRank, which includes a README and an example for running it. The ERDS library of Enamine compounds used in this work is available at our Zenodo data depository. We chose to provide a copy of this compound library as it is no longer available from Enamine, having been superseded by a 95.6M molecule library (that should contain the 67.5M subset we use). The Zenodo data depository also contains the ranks for all binders and decoys for each of the DUD-E targets, to permit others to perform detailed comparisons as we were able to do only because the team behind Glide chose to make the data public. We think this should be the norm and not the exception. The Zenodo data depository also contains the binding-site definitions used for each DUD-E target and all supplementary tables. This depository can be found at https://doi.org/10.5281/zenodo.16861024. NRGRank can also be used as part of the NRGSuite-Qt ^40^ to screen pre-loaded compound databases including FDA approved drugs, all ligands present in the PDB and all tetrapeptides. Additionally, the NRGSuite-Qt offers the possibility for users to load their own ligand datasets in SMILE format. The NRGSuite-Qt PyMOL plugin also includes GetCleft ^29^ for the detection and refinement of cavities/binding-sites, FlexAID for re-scoring and pose determination, NRGTEN ^41^ for the generation of target conformational ensembles and IsoMIF ^42^ for the detection of binding-site similarities, particularly useful when screening ligands are present in the PDB ^43^, allowing one to determine if the high-ranking screening and docking scores reflect common interactions ^43^.

## 3 Conclusions

NRGRank is a structurally-informed virtual screening Python package capable of ultra-massive virtual screening. Its scoring function is based on the exhaustive quantification of pairwise atomic interaction energies using 40 atom types. NRGRank shows comparable performance in virtual screening experiments to docking software Glide, AutoDock Vina and DOCK 3.7, particularly for targets in apo form and AF2 models, while being 1-to-2 orders of magnitude faster. A re-scoring experiment with Glide demonstrates that re-scoring can identify additional binders. The re-scoring process enriches the top-ranked results by recovering binders that were initially missed by NRGRank, while still retaining those binders uniquely identified by NRGRank. However, even without re-scoring, NRGRank finds a distinct and substantial subset of binders compared to Glide. We conjecture that the same is true for any pair of virtual screening software. That is, that each has unique biases that lead to the detection of different sets of binders. Therefore, instead of seeking to use different virtual screening software to validate results, users are more likely to benefit from the union of top results from various software rather than the intersection of top results. We show that while NRGRank produces screening artifacts, they can be eliminated by focusing on target-specific top-scoring compounds, which are found for every one of the 102 DUD-E targets we screened the 65-million Enamine REAL Diversity Subset (ERDS) against. We show that NRGRank can discriminate known binders from the large library ERDS background and that re-scoring can help judge if a molecule represents a screening artifact or a candidate for further analysis. Lastly, we show that NRGRank detects different top-ranking molecules even for closely related targets, namely individual members of the PIM kinase family. The simplicity of NRGRank, which does not require expert pre-processing or target refinement, combined with its minimal computational resource requirements, equivalent accuracy for apo form or model targets, complementary results to docking-based screening software, and high speed, makes it a valuable addition to the arsenal of computational tools for ultra-massive virtual screening with low barriers to adoption.

## 4 Aknowledgements

We thank the Digital Research Alliance of Canada for providing computational resources. RJN is a member of the Quebec Network for Research on Protein Function, Engineering, and Applications (PROTEO).

## 5 Methods

Three inputs are necessary to run NRGRank: Target structure, binding-site definition and ligand structures. The target structure and binding-site definition are used in a pre-processing step to create a lookup grid of pre-calculated pairwise atom type interactions. This pre-processed grid and the ligand structures are then used by NRGRank to probe exhaustively each ligand rotation and translation within the binding-site. The target and ligands are required to be in Mol2 format with SYBYL atom types assigned.

### 5.1 TARGET AND LIGAND PREPARATION

NRGRank utilizes a scoring function based on SYBYL atom types. As such, it is necessary to map both protein and ligand atoms into SYBYL atom types. The Mol2 structure file format automatically assigns SYBYL atom types to both protein and ligand atoms. The first step in preparing the target is converting the original protein structure and ligand files to the Mol2 format. There are many ways to do this, for example using PyMOL ^44^ or Open Babel ^45^. RDKit ^46^ was used to generate one conformer for each ligand from SMILES. We used the ETKDGv3 conformer generator in EmbedMultipleConfs from the rdkit.Chem.AllChem module.

### 5.2 BINDING-SITE DEFINITION

Binding-sites were defined as previously ^29, 47^ using the GetCleft algorithm, which is a C programming language implementation of the Surfnet algorithm ^48^. Briefly, the algorithm defines a binding-site as the largest set of intersecting spheres that can be placed at the midpoint between any pair of atoms without overlapping with the van der Walls radii of any protein atom and with sphere radii within lower (rl) and upper (ru) values. Careful choice of the lower and upper values permits to fine-tune the shape of the binding-site volume without “spilling over” across cavities that ought to be considered unconnected by visual inspection. In cases where adjustments were necessary, we fine-tuned the sphere radii away from default values. Targets in the DUD-E dataset are provided in the holo form. In these cases, the GetCleft cavity containing the bound ligand was selected for the experiments using rl and ru 1.75 Å and 3.75 Å respectively. For experiments with apo and AF2 structures, we utilized the structures provided by the authors ^10^ and the equivalent GetCleft cavity. That is, the cavity in the apo or AF2 structure that overlaps with the ligand-bound cavity in the holo form when the two protein structures are superimposed. It is worth mentioning that we do not constraint the definition of the binding-site volume in any way with the bound ligand in the holo form, a practice often employed in the benchmarking of docking algorithms and that greatly biases the analysis by severely restricting the volume of search space. As NRGRank performs an exhaustive search, the additional search space at most could impact execution time. In the case of PIM1, PIM2 and PIM3, we selected the binding pocket with the largest volume, that coincides with the ATP binding pocket.

### 5.3 TARGET PRE-PROCESSING

A pre-processing step is required to create a lookup grid of pseudo-energies for each atom type based on the distribution of protein atoms in the structure. The Mol2 protein structure file is loaded into the software and atoms are indexed into a grid of index-cubes. As NRGRank is somewhat insensitive to the choice of index-cube width (Supplementary Figure S1), we chose a width with a physical explanation, namely equal to two times the maximum van der Waals atom radius (rvdW) of all Sybyl atom types (1.88 Å) added to two times the rvdW of water (1.44 Å), for a total of 6.56 Å. This represents the distance between the center of the two biggest Sybyl atom types separated by a water molecule. s such, an index cube may contain two atoms in contact, as defined by Surfaces ^49^ and FlexAID ^29^. Subsequently, for each of these cubes, the energy matrix developed for FlexAID is used to calculate the pseudo-energy of each possible Sybyl atom type with all protein atoms within the central and adjacent cubes, for a total of 27 cubes. The binding energy for one Sybyl atom type is given by:

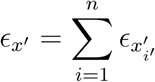

where *ɛ_x′_* is the total energy for an atom *x* of Sybyl atom type *x^′^* located in a cube and all atoms located within 1 cube excluding other ligand atoms. The pairwise interaction energy between atom *x* and atom *i* of Sybyl atom type *i^′^*is given by *ɛ_x′i′_*. Therefore, a ligand atom of a particular atom-type will contribute an amount *ɛ_x′_* irrespective of where the protein atoms that contributes to that interaction is located within a 7622.11 Å^3^ volume (equal to the volume of 27 cubes with 6.56 Å side). As a result, NRGRank is insensitive to ligand conformers, side-chain rotamers and minor backbone movements, any movements that maintain the atoms within the same interaction volume contribute equally.

The binding site volume provided as input is utilized to generate ligand anchor points as a grid spaced at intervals of 1.5 Å. Each anchor point represents a position at which each ligand will be tested. The center of geometry of each ligand is placed on each anchor point in the binding site. For each anchor point, the ligand is rotated 9 times per axis to sample 729 different orientations. The binding energy of a pose is the cumulative sum of the binding energies associated with each ligand atom, as calculated during the pre-processing phase. The final score for a compound corresponds to the lowest pseudo-energy obtained in the exhaustive search. Given the simplified nature of the scoring, the pose associated to that score is not saved, only the scores and relative ranking of all molecules tested.

### 5.4 EVALUATION OF PERFORMANCE ON APO PROTEIN STRUCTURES AND ALPHAFOLD MODELS

We utilized the apo, holo, and AF2 structures on a subset of 37 DUD-E targets as provided by the authors for comparisons with Glide ^10^. The structures provided in maegz format were converted to mol2 files utilizing Schrödinger Maestro (Release 2024-2; Schrödinger, LLC, New York, NY, 2024). The methodology outlined in the binding site definition section was used for holo structures. To mimic the binding site preparation for Glide, the holo binding site definition was used for apo and AlphaFold 2 structures.

When calculating ligand percentile rank from Glide, the best scoring result for each molecule was retained, and duplicate or worse scores were discarded.

### 5.5 EVALUATION OF PERFORMANCE ON HOLO PROTEIN STRUCTURE

For each of the 37 targets used in the Glide paper, the crystal structure was retrieved from the DUD-E database in the pdb format and converted to the Mol2 format using PyMOL. Next, binding sites were defined around crystallized ligands using GetCleft with an rl of 1.75 Å and ru equal to 3.75 Å or adjusted when the ligand did not fit within the binding site. Target structures and ligand Mol2 files from the DUD-E website were then preprocessed using NRGRank. Subsequently, NRGRank screening was ran on the Narval cluster managed by Calcul Québec (calculquebec.ca) and the Digital Research Alliance of Canada (alliance-can.ca). This required approximately 30 minutes for the 102 targets in DUD-E. Finally, EF1 was calculated as described by Jain et al. ^50^.

### 5.6 EVALUATION OF NON-PROPERTY MATCHED DECOYS

All 102 targets were obtained from the DUD-E dataset. Known binders were obtained from ChEMBL ^51, 52^ and respect the following criteria: EC_50_, K*_i_*, K*_d_* values *≤* 1 *µ*M. The decoys are a set of diverse molecules comprising 67.5 million compounds from the Enamine REAL diversity subset. This resulted in the evaluation of over 6.7 billion molecules. The ERDS 67.5M compounds dataset and the list of ChEMBL known binders for DUD-E targets are available online at the Zenodo data depository.

## Supplementary Figures

**Figure S1:**
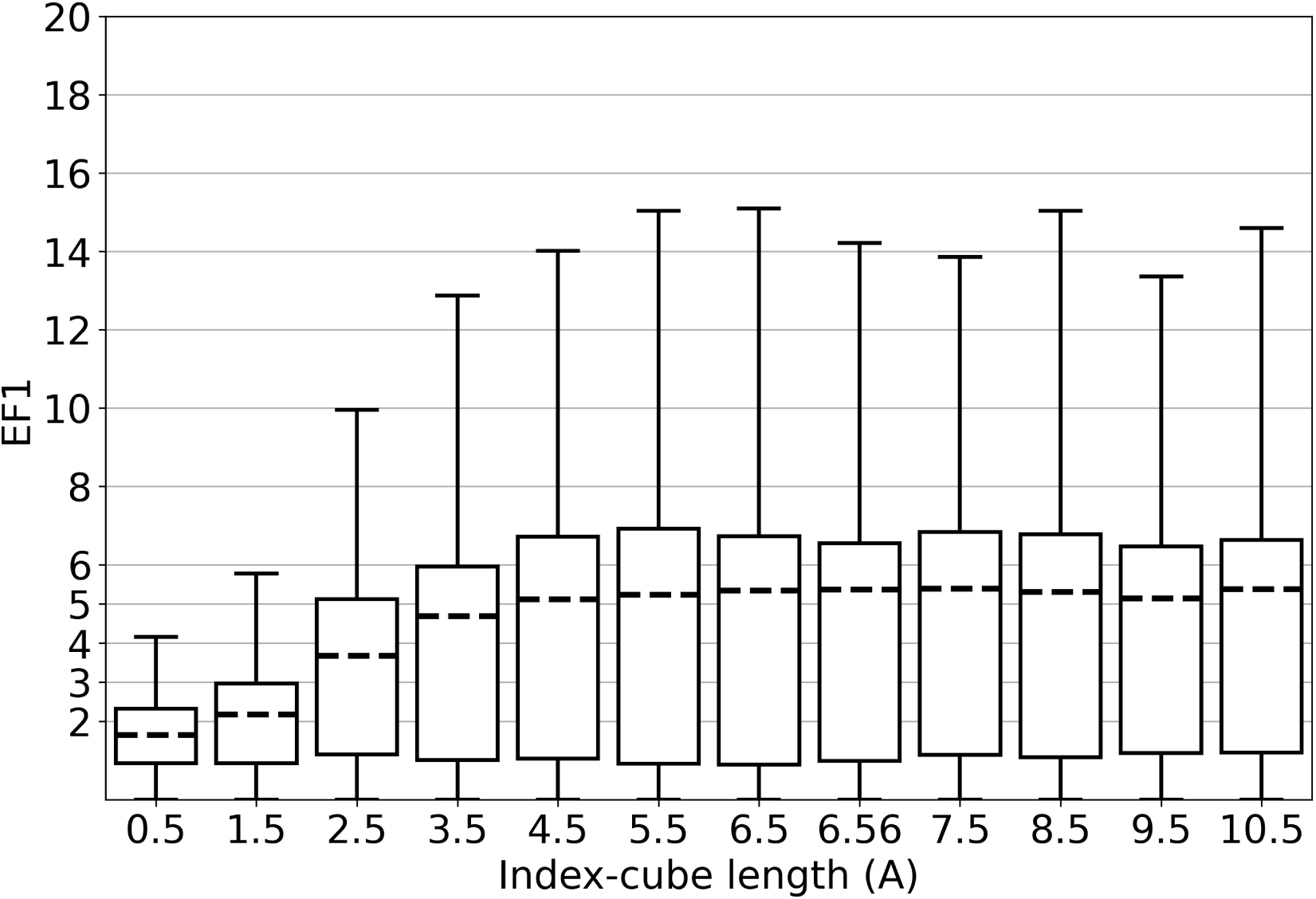
Effect of index-cube length on EF1.

**Figure S2:**
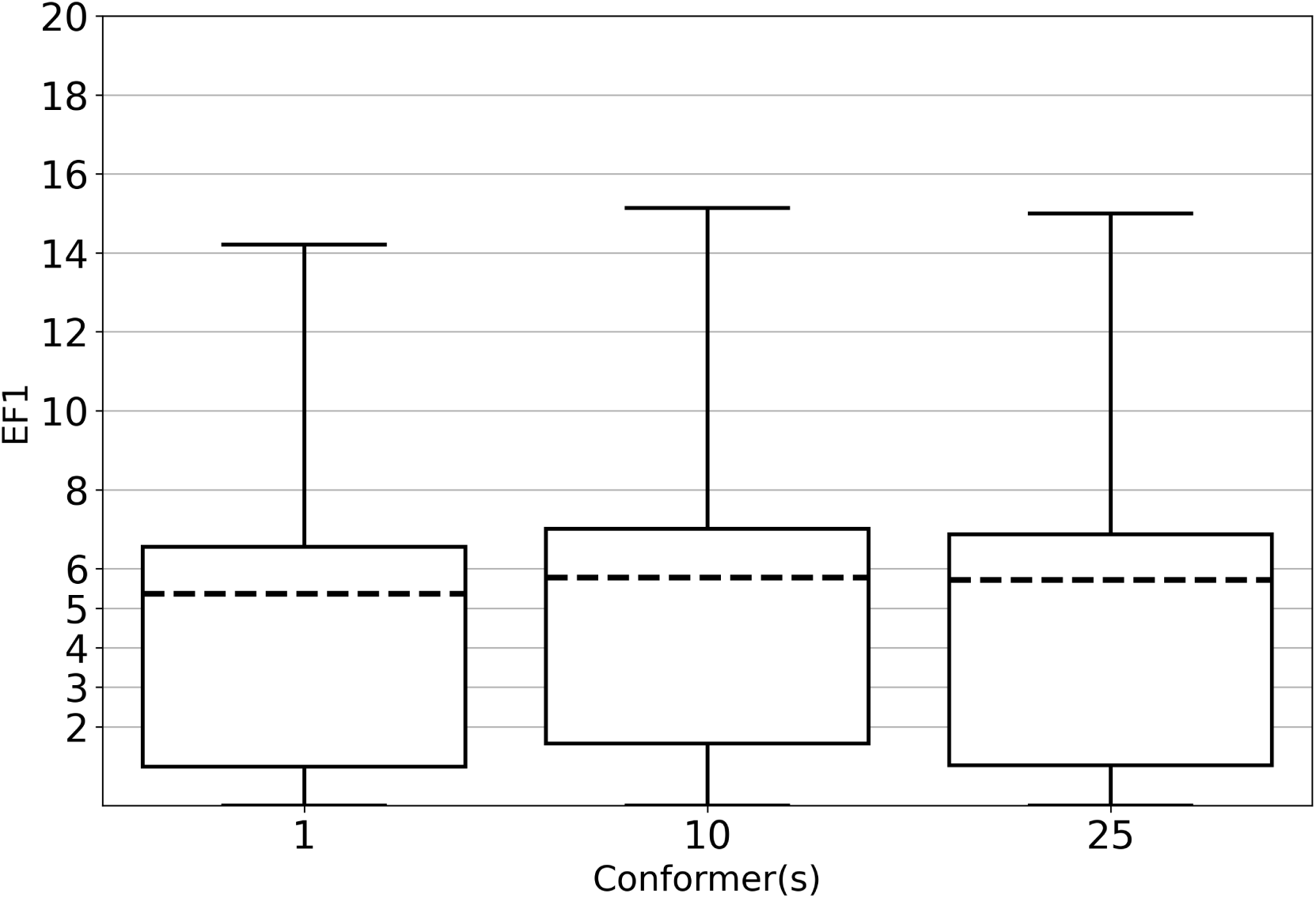
Effect of number of conformers used on EF1.

**Figure S3:**
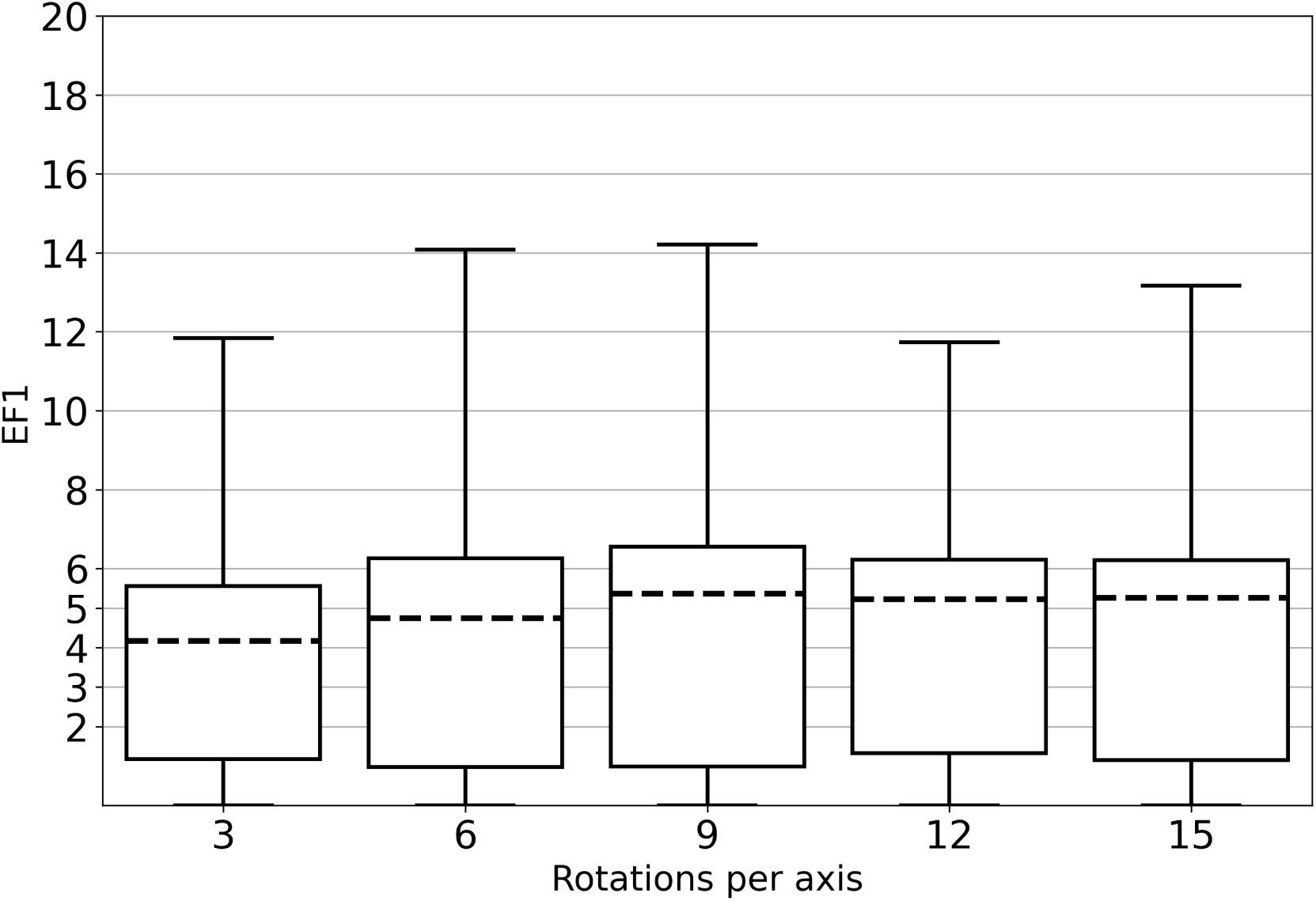
Effect of number of rotations per axis on EF1.

**Figure S4:**
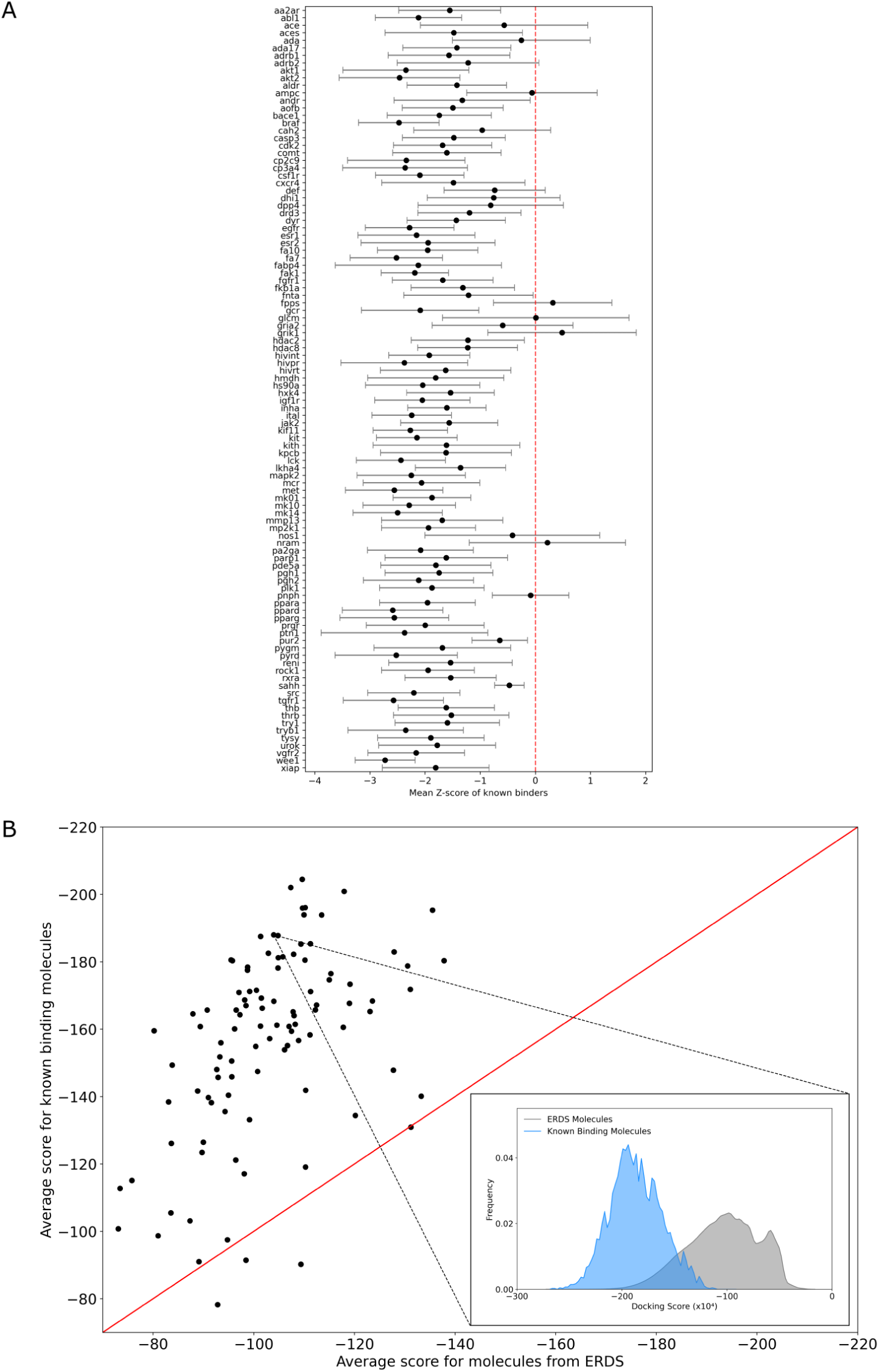
Ranking ERDS (Enamine REAL Diversity subset) on 102 targets of the DUD-E dataset. Average ranking score for known binders and the average for the ERDS compounds for each target (the more negative the better the score is). The list of ERDS and known binder distribution averages as well as number of known binders can be found in Supplementary Table S2.

**Figure S5:**
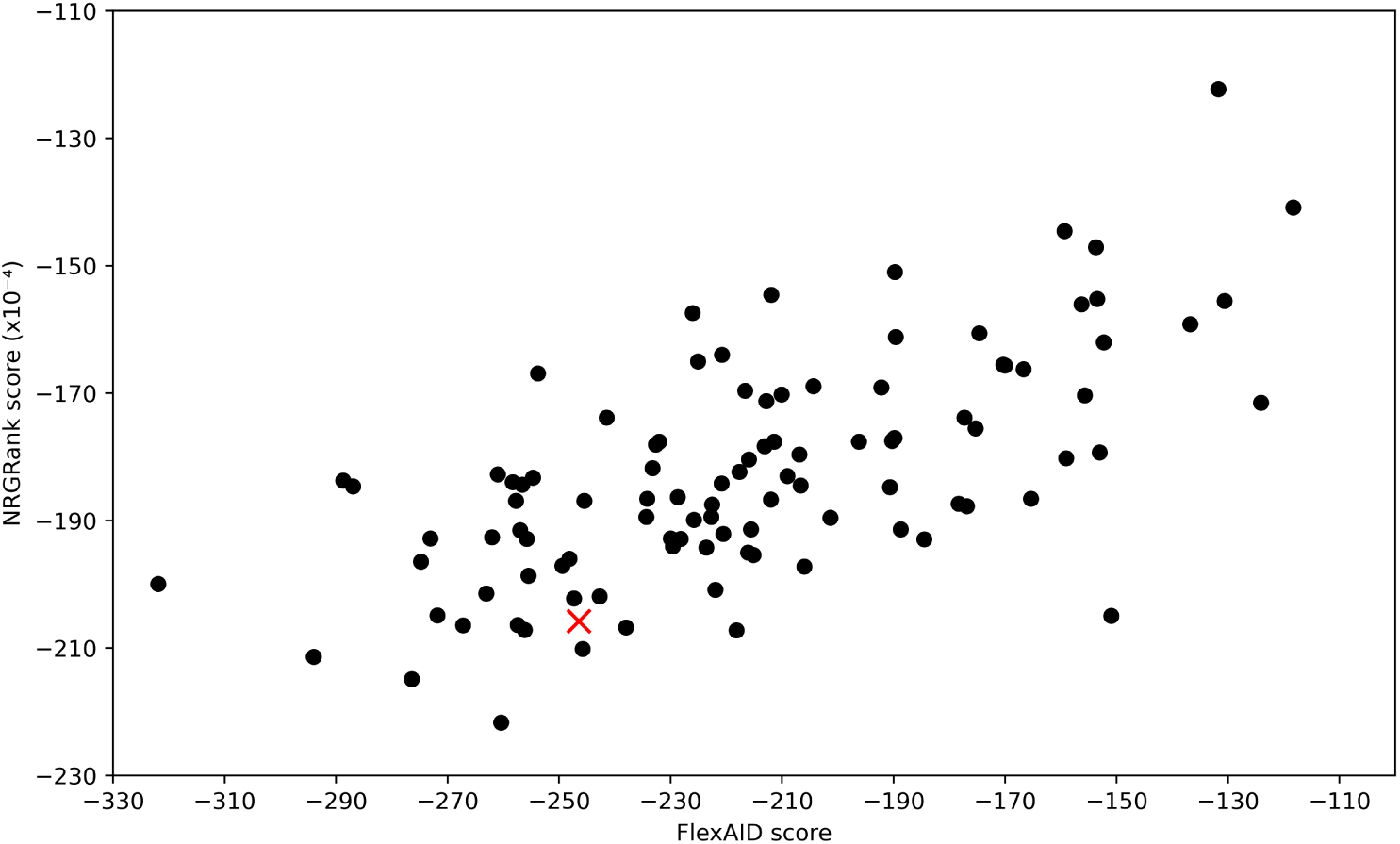
Comparison of FlexAID and NRGRank scores for molecule S_994 across 102 DUD-E targets. The target *sahh* is indicated with a red cross. A Pearson correlation coefficient of 0.65 was observed. FlexAID scores had a mean of *−*215.75 *±* 41.61 AU, while NRGRank scores had a mean of (*−*182.77 *±* 17.57) *×* 10^4^.

**Figure S6:**
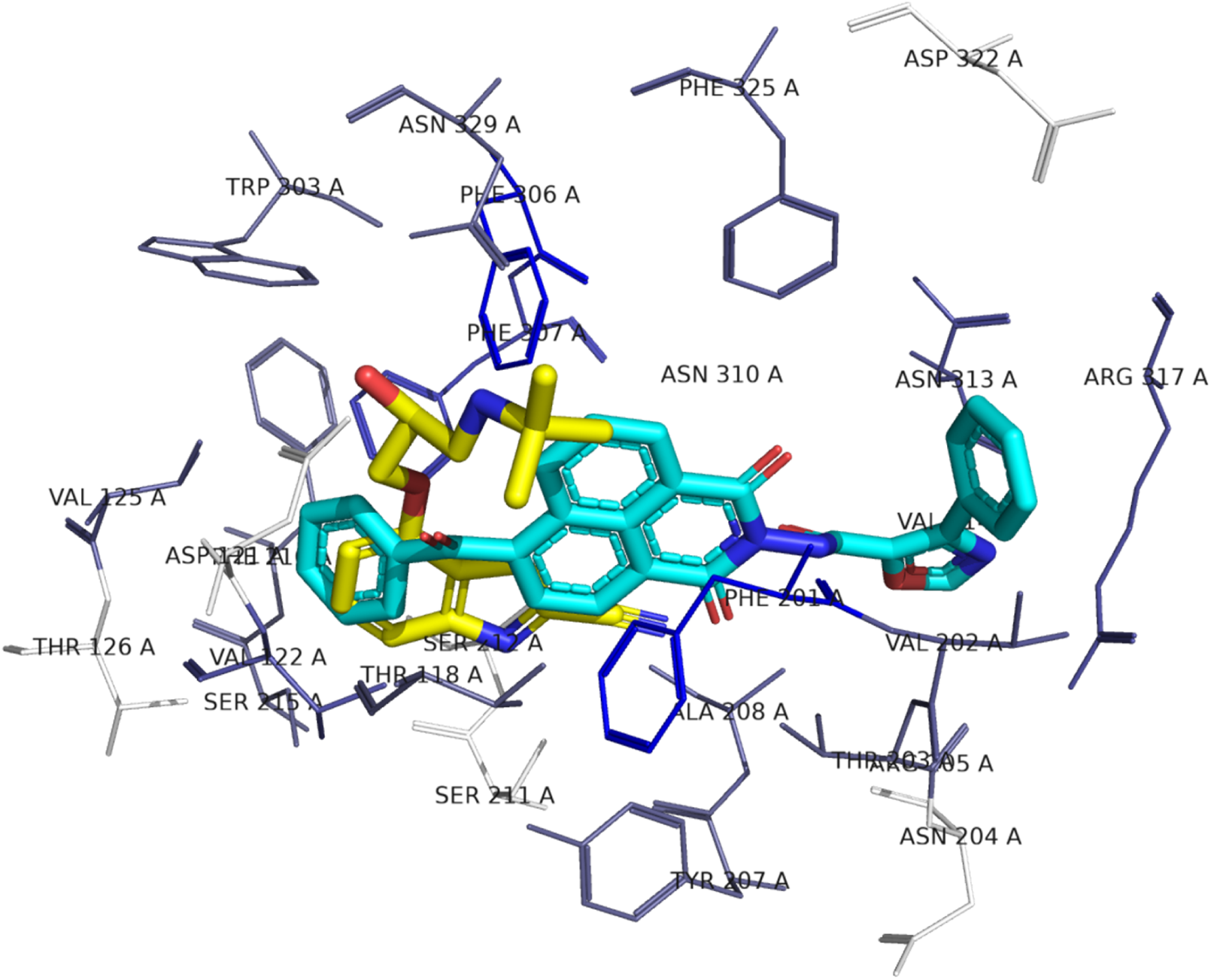
FlexAID docking pose of ERDS compound. *S***_**652 **on adrb1.** The ERDS compound *s*_22 22609224 10640652 (*S*_652 in cyan), found as the top-ranking ERDS compound by NRGRank, was docked with FlexAID. Binding site side-chains are colored in a white to blue scale. The brighter the shade of blue, the more that residue contributes to the interaction with *S*_652.

**Figure S7:**
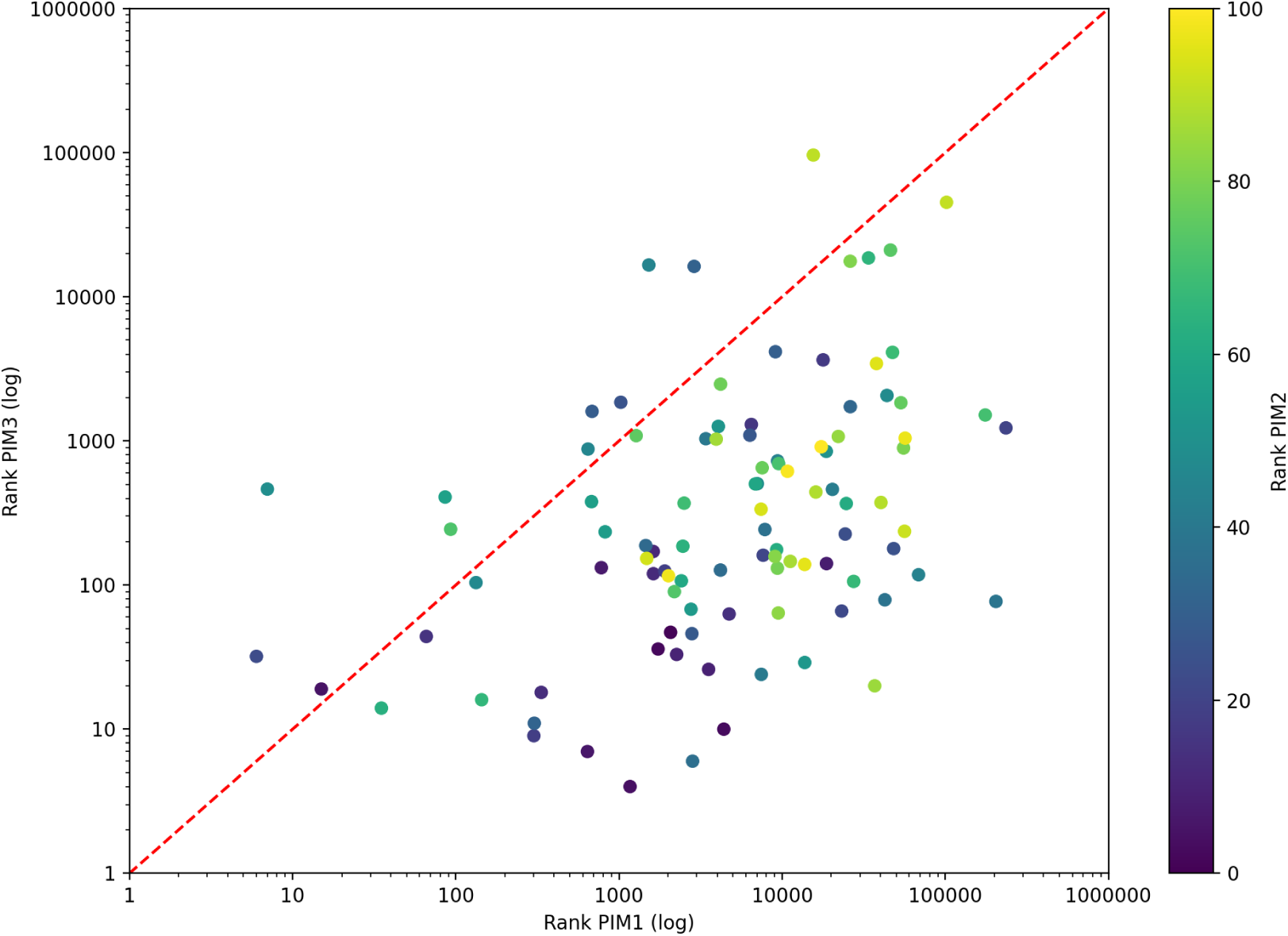
Rank comparison on PIM1 and PIM3 for the top 100 molecules ranked for PIM2. Among molecules in the top 100, a number have poor ranks (10,000 and above) particularly for PIM1 but also for PIM3, in agreement with the fact that PIM2 has less mutations with respect to PIM3 than to PIM1.

**Figure S8:**
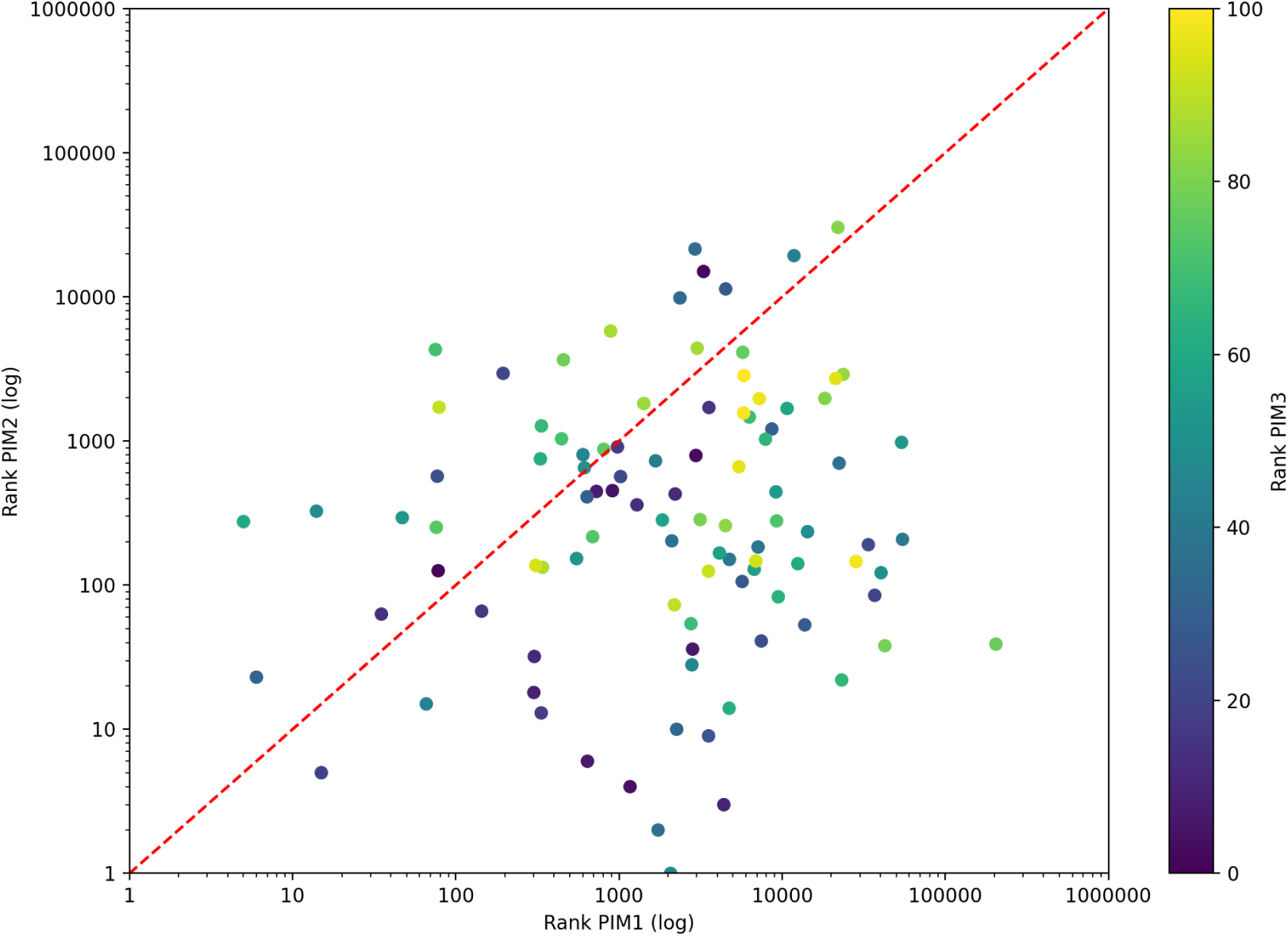
Rank comparison on PIM1 and PIM2 for the top 100 molecules ranked for PIM3. Among molecules in the top 100, a number have poor ranks (10,000 and above) particularly for PIM1 but also some for PIM2. While PIM3 has 4 mutations compared to PIM2 and only 3 compared to PIM1, it has only one mutation for residues whose side-chains face the binding-site cavity. The cavity side-chain facing mutation from PIM3 to PIM1 is an ALA to VAL, a more conservative mutation than that between PIM3 and PIM2 where the alteration is a GLU to LEU. This is reflected on the fact that ranks for the top PIM3 molecules on PIM1 is shifted to the left when compared to the ranks of the top 100 molecules for PIM2 on PIM1 (Supplementary Figure S5).

**Supplementary Table S1:** Information about targets and EF1 for various methods on the DUD-E dataset. Supplementary table 1 is available in the Zenodo data depository as s1.csv

**Supplementary Table S2:**
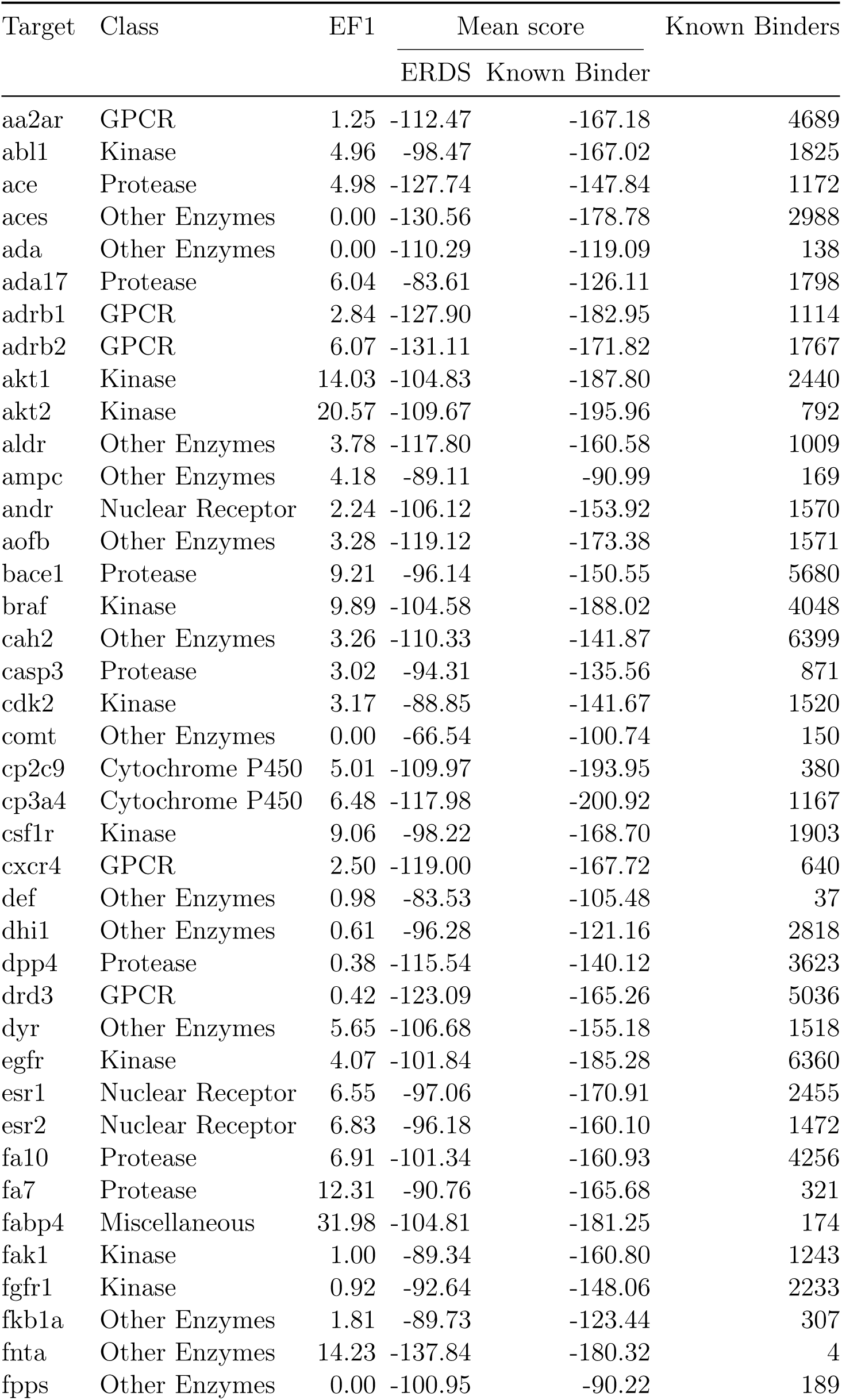
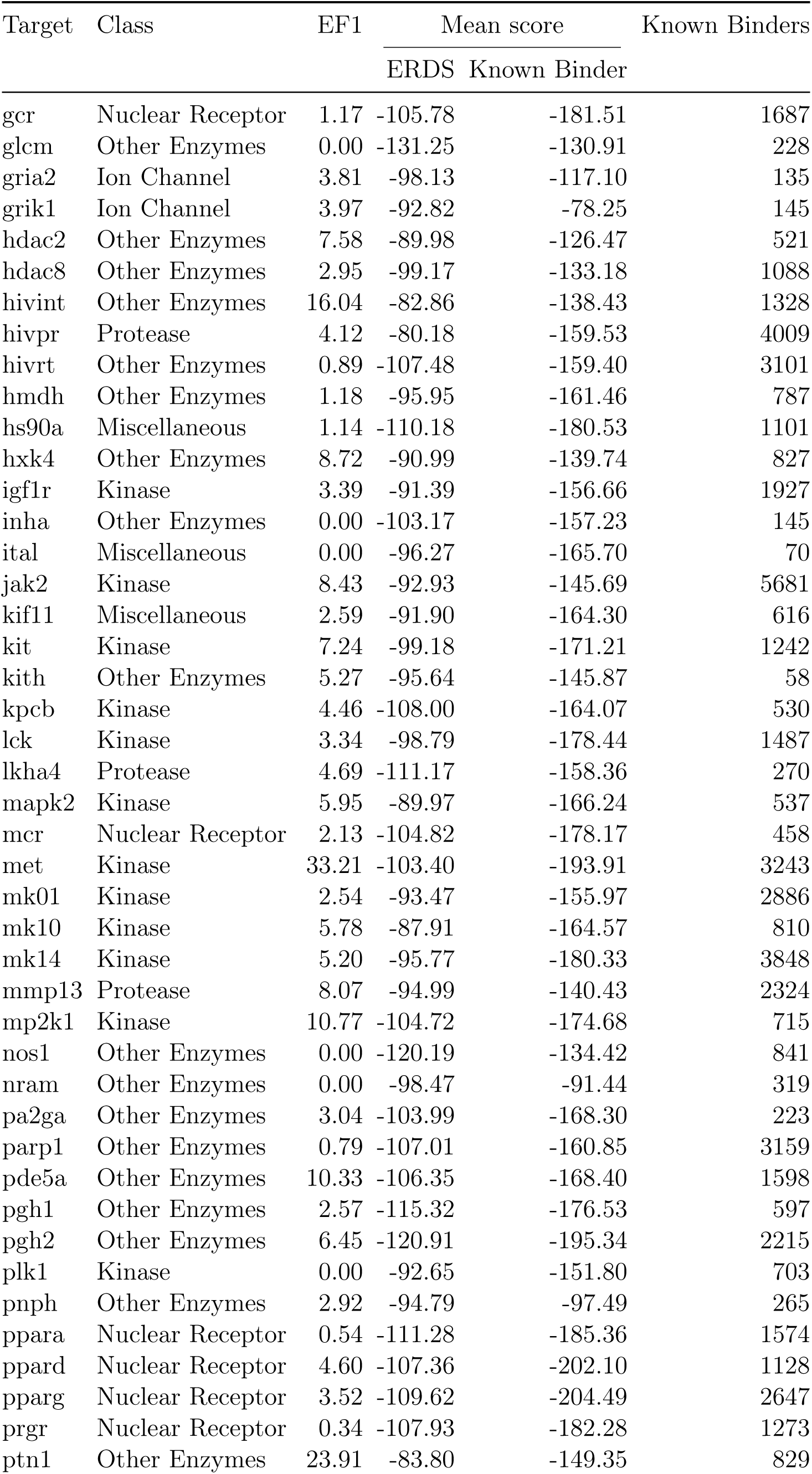
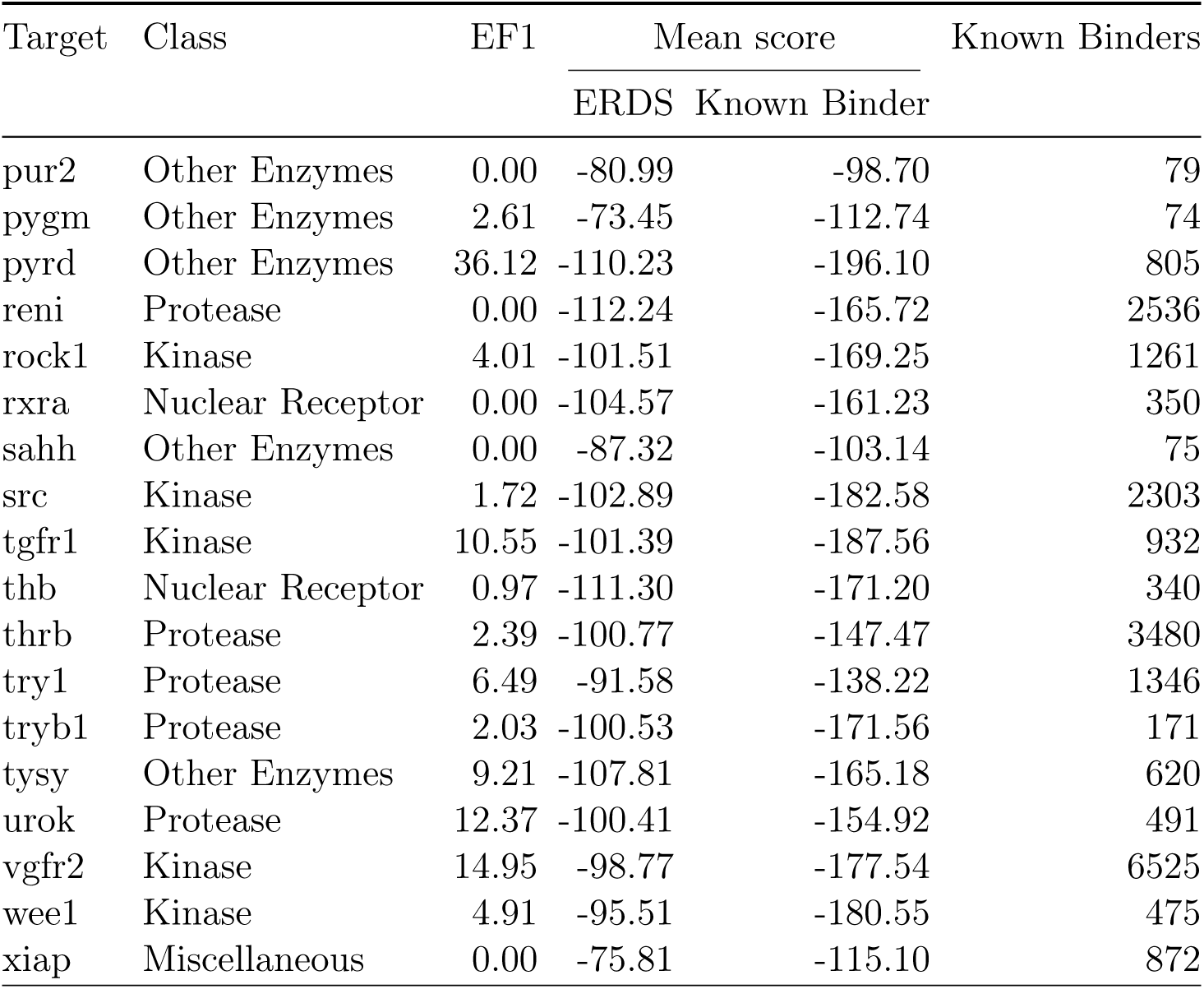
ERDS and known binder mean ranks with EF1 and binder counts. Average ranking score (mean) for ERDS compounds and known binders for each target, along with EF1 and the number of screened known binders.

**Supplementary Table S3:**
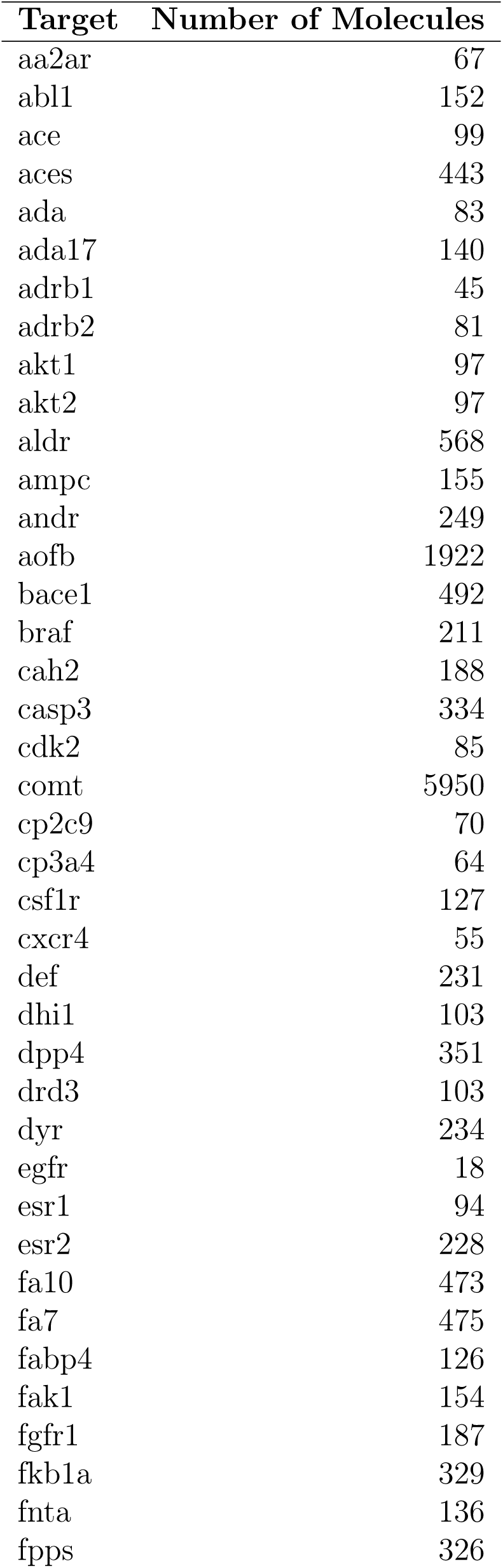
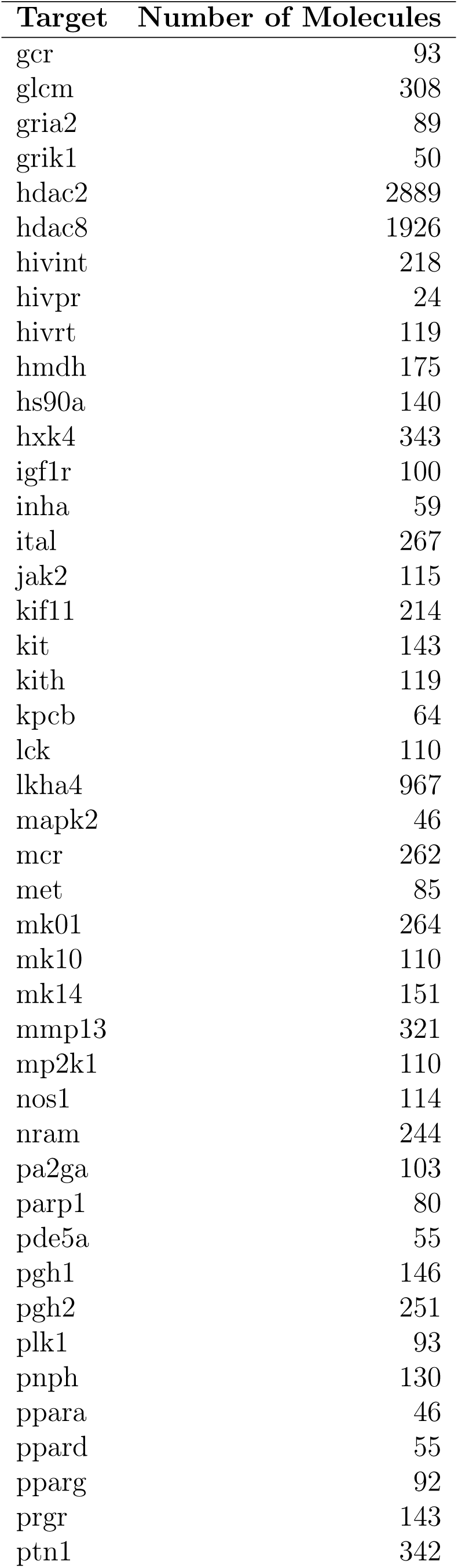
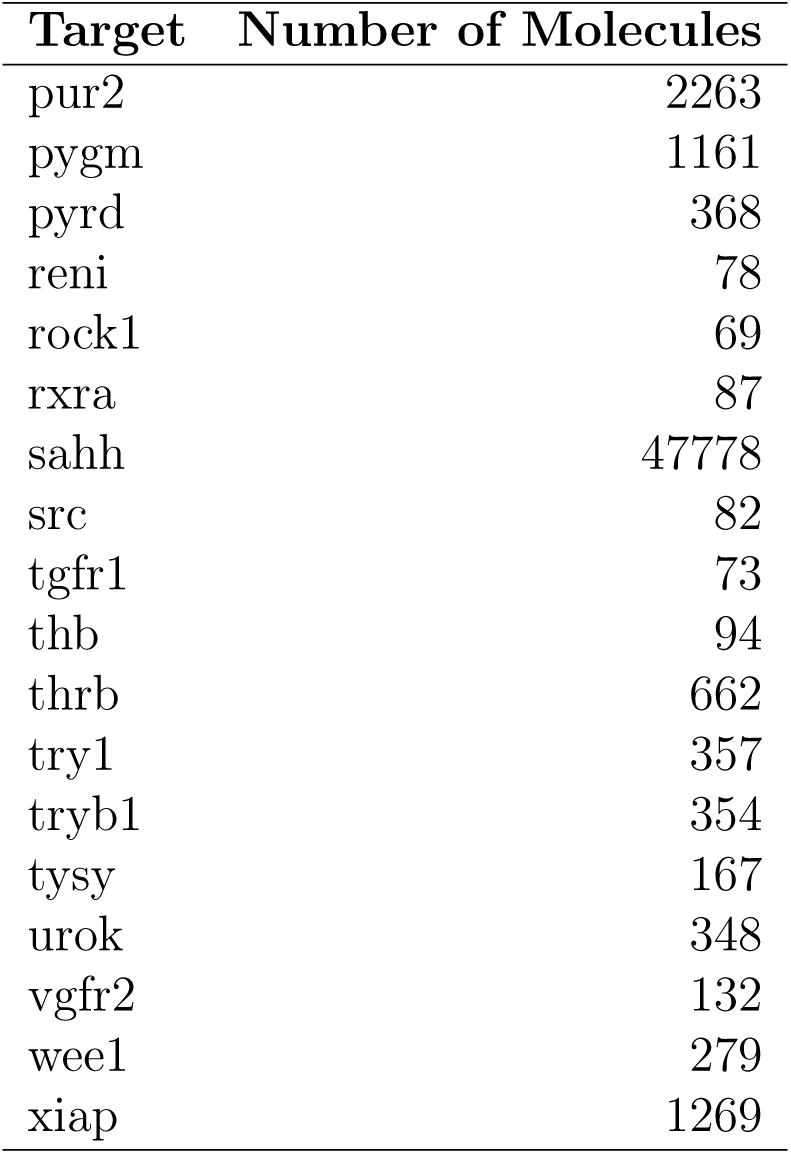
Number of molecules with pProp *≥* 3 and ΔpProp > 2 per target.

## References

1. Michael, S. et al. A robotic platform for quantitative high-throughput screening. eng. Assay and drug development technologies 6, 637–657 (Oct. 2008).

2. Grygorenko, O. O. et al. Generating Multibillion Chemical Space of Readily Accessible Screening Compounds. eng. iScience 23, 101681 (Oct. 2020).

3. Lyu, J. et al. Ultra-large library docking for discovering new chemotypes. eng. Nature 566, 224–229 (Feb. 2019).

4. Lyu, J. et al. Modeling the expansion of virtual screening libraries. eng. Nature chemical biology 19, 712–718 (June 2023).

5. Liu, F. et al. The impact of library size and scale of testing on virtual screening. Nature Chemical Biology 21, 1039–1045 (Jan. 2025).

6. Singh, I. et al. Structure-Based Discovery of Inhibitors of the SARS-CoV-2 Nsp14 N7-Methyltransferase. Journal of Medicinal Chemistry 66, 7785–7803 (June 2023).

7. Gahbauer, S. et al. Docking for EP4R antagonists active against inflammatory pain. eng. Nature communications 14, 8067 (Dec. 2023).

8. Kelleher, K. J. et al. Pharos 2023: an integrated resource for the understudied human proteome. eng. Nucleic acids research 51, D1405–D1416 (Jan. 2023).

9. Jumper, J. et al. Highly accurate protein structure prediction with AlphaFold. eng. Nature 596, 583–589 (Aug. 2021).

10. Zhang, Y. et al. Benchmarking Refined and Unrefined AlphaFold2 Structures for Hit Discovery. Journal of Chemical Information and Modeling 63, 1656–1667 (Mar. 2023).

11. Friesner, R. A. et al. Glide: A New Approach for Rapid, Accurate Docking and Scoring. 1. Method and Assessment of Docking Accuracy. Journal of Medicinal Chemistry 47, 1739–1749 (Mar. 2004).

12. Mysinger, M. M. et al. Directory of useful decoys, enhanced (DUD-E): better ligands and decoys for better benchmarking. eng. Journal of medicinal chemistry 55, 6582– 6594 (July 2012).

13. Sabe, V. T. et al. Current trends in computer aided drug design and a highlight of drugs discovered via computational techniques: A review. European Journal of Medicinal Chemistry 224, 113705 (2021).

14. Trott, O. et al. AutoDock Vina: improving the speed and accuracy of docking with a new scoring function, efficient optimization, and multithreading. eng. Journal of computational chemistry 31, 455–461 (Jan. 2010).

15. Eberhardt, J. et al. AutoDock Vina 1.2.0: New Docking Methods, Expanded Force Field, and Python Bindings. eng. Journal of chemical information and modeling 61, 3891–3898 (Aug. 2021).

16. Balius, T. E. et al. DOCK 6: Incorporating hierarchical traversal through precomputed ligand conformations to enable large-scale docking. Journal of Computational Chemistry 45, 47–63 (Jan. 2024).

17. McGann, M. FRED Pose Prediction and Virtual Screening Accuracy. Journal of Chemical Information and Modeling 51, 578–596 (2011).

18. Diller, D. J. et al. High throughput docking for library design and library prioritization. Proteins: Structure, Function, and Genetics 43, 113–124 (2001).

19. Arul Murugan, N., et al. Artificial intelligence in virtual screening: Models versus experiments. Drug Discovery Today 27, 1913–1923 (2022).

20. Zheng, L., et al. OnionNet: a Multiple-Layer Intermolecular-Contact-Based Convolutional Neural Network for Protein-Ligand Binding Affinity Prediction. eng. ACS omega 4, 15956–15965 (Sept. 2019).

21. Buttenschoen, M. et al. PoseBusters: AI-based docking methods fail to generate physically valid poses or generalise to novel sequences. eng. Chemical science 15, 3130–3139 (Dec. 2023).

22. Chen, L. et al. Hidden bias in the DUD-E dataset leads to misleading performance of deep learning in structure-based virtual screening. eng. PloS one 14, e0220113– e0220113 (Aug. 2019).

23. Yu, Y. et al. Uni-Dock: GPU-Accelerated Docking Enables Ultralarge Virtual Screening. Journal of Chemical Theory and Computation 19, 3336–3345 (2023).

24. Ding, J. et al. Vina-GPU 2.0: Further Accelerating AutoDock Vina and Its Derivatives with Graphics Processing Units. Journal of Chemical Information and Modeling 63, 1982–1998 (Apr. 2023).

25. Gaudreault, F. et al. Side-chain rotamer changes upon ligand binding: common, crucial, correlate with entropy and rearrange hydrogen bonding. eng. *Bioinformatics (Oxford*, England) 28, i423–i430 (Sept. 2012).

26. Xu, M. et al. Systematic Investigation of Docking Failures in Large-Scale Structure-Based Virtual Screening. eng. ACS omega 7, 39417–39428 (Oct. 2022).

27. Baell, J. B. et al. Seven Year Itch: Pan-Assay Interference Compounds (PAINS) in 2017—Utility and Limitations. ACS Chemical Biology 13, 36–44 (Dec. 2017).

28. Narayanan, S. R. et al. 9-(Trans-2’-trans-3’-dihydroxycyclopent-4’-enyl) derivatives of adenine and 3-deazaadenine: potent inhibitors of bovine liver S-adenosylhomocysteine hydrolase. Journal of Medicinal Chemistry 31, 500–503 (Mar. 1988).

29. Gaudreault, F. et al. FlexAID: Revisiting Docking on Non-Native-Complex Structures. Journal of Chemical Information and Modeling 55, 1323–1336 (2015).

30. Yang, X. et al. Catalytic Strategy of S-Adenosyl-l-homocysteine Hydrolase: Transition-State Stabilization and the Avoidance of Abortive Reactions. Biochemistry 42, 1900– 1909 (Jan. 2003).

31. Engel, G. et al. (+/-)[125Iodo]cyanopindolol, a new ligand for ?-adrenoceptors: Identification and quantitation of subclasses of ?-adrenoceptors in guinea pig. Naunyn-Schmiedeberg’s Archives of Pharmacology 317, 277–285 (1981).

32. Warne, T. et al. Structure of a beta1-adrenergic G-protein-coupled receptor. eng. Nature 454, 486–491 (July 2008).

33. Speers, C. et al. Identification of novel kinase targets for the treatment of estrogen receptor-negative breast cancer. eng. Clinical cancer research : an official journal of the American Association for Cancer Research 15, 6327–6340 (Oct. 2009).

34. Bianchini, G. et al. Prognostic and Therapeutic Implications of Distinct Kinase Expression Patterns in Different Subtypes of Breast Cancer. Cancer Research 70, 8852–8862 (2010).

35. Mahata, S. et al. PIM1/STAT3 axis: a potential co-targeted therapeutic approach in triple-negative breast cancer. Medical Oncology 39 (2022).

36. Yuan, Y. et al. PIM1 promotes hepatic conversion by suppressing reprogramming-induced ferroptosis and cell cycle arrest. Nature Communications 13, 5237 (Sept. 2022).

37. Zhao, W. et al. PIM1: a promising target in patients with triple-negative breast cancer. Medical Oncology 34 (2017).

38. Brasó-Maristany, F. et al. PIM1 kinase regulates cell death, tumor growth and chemotherapy response in triple-negative breast cancer. eng. Nature medicine 22, 1303–1313 (Nov. 2016).

39. Merkel, A. L. et al. PIM1 kinase as a target for cancer therapy. Expert Opinion on Investigational Drugs 21, 425–436 (Apr. 2012).

40. Galdino, G. T. et al. NRGSuite-Qt: a PyMOL plugin for high-throughput virtual screening, molecular docking, normal-mode analysis, the study of molecular interactions, and the detection of binding-site similarities. Bioinformatics Advances 5 (ed Lengauer, T.) (Dec. 2024).

41. Mailhot, O. et al. The NRGTEN Python package: an extensible toolkit for coarse-grained normal mode analysis of proteins, nucleic acids, small molecules and their complexes. Bioinformatics 37, 3369–3371 (2021).

42. Chartier, M. et al. Detection of Binding Site Molecular Interaction Field Similarities. Journal of Chemical Information and Modeling 55, 1600–1615 (2015).

43. Rodríguez-Salazar, C. A. et al. Ebola virus VP35 interacts non-covalently with ubiquitin chains to promote viral replication. eng. PLoS biology 22, e3002544–e3002544 (Feb. 2024).

44. DeLano, W. The PyMOL Molecular Graphics System

45. O’Boyle, N. M. et al. Open Babel: An open chemical toolbox. eng. Journal of cheminformatics 3, 33 (Oct. 2011).

46. Riniker, S. et al. Better Informed Distance Geometry: Using What We Know To Improve Conformation Generation. Journal of Chemical Information and Modeling 55, 2562– 2574 (2015).

47. Gaudreault, F. et al. NRGsuite: a PyMOL plugin to perform docking simulations in real time using FlexAID. eng. Bioinformatics (Oxford, England) 31, 3856–3858 (Dec. 2015).

48. Laskowski, R. A. SURFNET: A program for visualizing molecular surfaces, cavities, and intermolecular interactions. Journal of Molecular Graphics 13, 323–330 (1995).

49. Teruel, N. et al. Surfaces: a software to quantify and visualize interactions within and between proteins and ligands. Bioinformatics 39 (Oct. 2023).

50. Jain, A. N. Bias, reporting, and sharing: computational evaluations of docking methods. Journal of Computer-Aided Molecular Design 22, 201–212 (2007).

51. Davies, M. et al. ChEMBL web services: streamlining access to drug discovery data and utilities. eng. Nucleic acids research 43, W612–W620 (July 2015).

52. Zdrazil, B. et al. The ChEMBL Database in 2023: a drug discovery platform spanning multiple bioactivity data types and time periods. eng. Nucleic acids research 52, D1180–D1192 (Jan. 2024).

